# Substantial Downregulation of Mitochondrial and Peroxisomal Proteins during Acute Kidney Injury revealed by Data-Independent Acquisition Proteomics

**DOI:** 10.1101/2023.02.26.530107

**Authors:** Jordan B. Burton, Anne Silva-Barbosa, Joanna Bons, Jacob Rose, Katherine Pfister, Fabia Simona, Tejas Gandhi, Lukas Reiter, Oliver Bernhardt, Christie L. Hunter, Eric S Goetzman, Sunder Sims-Lucas, Birgit Schilling

**Author notes:** For correspondence Birgit Schilling, PhD, Buck Institute for Research on Aging, 8001 Redwood Boulevard, Novato, CA 94945, USA.

## Abstract

Acute kidney injury (AKI) manifests as a major health concern, particularly for the elderly. Understanding AKI-related proteome changes is critical for prevention and development of novel therapeutics to recover kidney function and to mitigate the susceptibility for recurrent AKI or development of chronic kidney disease. In this study, mouse kidneys were subjected to ischemia-reperfusion injury, and the contralateral kidneys remained uninjured to enable comparison and assess injury-induced changes in the kidney proteome. A fast-acquisition rate ZenoTOF 7600 mass spectrometer was introduced for data-independent acquisition (DIA) for comprehensive protein identification and quantification. Short microflow gradients and the generation of a deep kidney-specific spectral library allowed for high-throughput, comprehensive protein quantification. Upon AKI, the kidney proteome was completely remodeled, and over half of the 3,945 quantified protein groups changed significantly. Downregulated proteins in the injured kidney were involved in energy production, including numerous peroxisomal matrix proteins that function in fatty acid oxidation, such as ACOX1, CAT, EHHADH, ACOT4, ACOT8, and Scp2. Injured mice exhibited severely declined health. The comprehensive and sensitive kidney-specific DIA assays highlighted here feature high-throughput analytical capabilities to achieve deep coverage of the kidney proteome and will serve as useful tools for developing novel therapeutics to remediate kidney function.

## Introduction

Acute kidney injury (AKI) typically results in a sudden decrease in kidney function that occurs over a few hours or days. AKI increased 230% from 2000 to 2014, from 3.5 to 11.7 diagnoses and hospitalized patients per 1,000 individuals in the United States (1). The rise in the incidence of AKI is likely influenced by a growing aging population and the increased incidence of comorbidities, such as diabetes, hypertension, heart failure, and sepsis, that all affect a patient’s susceptibility to AKI (2, 3). Recurrent AKI within 6 months after the initial episode occurs in 25-31% of patients and is associated with increased risk of chronic kidney disease, renal failure and increased mortality (4–6). A meta-analysis of 312 studies, including 49,147,878 patients, primarily from high-income countries and nations with a total health expenditure ≥ 5% of the gross domestic product, found that AKI-associated mortality rates were as high as 23.9 ± 1.8% (7).

Kidney tissue is especially susceptible to damage due to the high metabolic rate needed for energy production to maintain kidney function (8–10). Kidney tissue is rich in mitochondria and peroxisomes, highly metabolically active, and relies on fatty acid β-oxidation (FAO) for energy production (10). Interestingly, parallel FAO metabolic pathways are present in the mitochondria and peroxisomes. Mitochondrial FAO is linked to the electron transport chain for energy production, which is suppressed after AKI. However, shifting to peroxisomal FAO may protect against the accumulation of long-chain fatty acids (10). Previously, our groups investigated the role of post-translational modifications, specifically succinylation, as a disease-modifying intervention for AKI investigating sirtuin 5 knockout mouse models (10). The knockdown of sirtuin 5, a desuccinylase, significantly increased protein succinylation in the injured kidneys. The latter improved the phenotypic response, and kidney health after AKI compared to the wild-type mice, which is likely the result of increased succinylation of proteins involved in fatty acid oxidation in both the mitochondria and peroxisomes (10). The current study primarily focuses on the proteome-wide changes of acutely injured kidney tissues after ischemic/reperfusion injury, compared to healthy contralateral kidney tissues in wild-type mice.

To comprehensively analyze and quantify kidney proteome profiles we chose a label-free, highly quantitative mass spectrometric methodology, referred to as data-independent acquisition (DIA). Briefly, mass spectrometric MS1 survey spectra are acquired of all peptide precursor ions present. Subsequently, tandem mass spectra (MS/MS) are acquired of segmented m/z windows that span the entire MS1 m/z precursor ion range (e.g., in this study, m/z 200–1500). Thus, data acquisition is independent of what actually was present in the MS1 precursor ion spectra, and all peptides are comprehensively sampled for MS/MS (11, 12). Typically, DIA-MS workflows result in complex MS/MS spectra as multiple peptides are present and fragmented in each window and then deciphered with sophisticated computational data-processing strategies (13). The use of variable windows m/z width for isolation of MS1 precursor ions to transmit peptide precursor ions for MS/MS fragmentation has improved quantification accuracy greatly in reducing complexity by using smaller DIA window width in MS1 m/z ranges that are particularly populated with precursor ions, such as m/z 500-700, and allowing for larger DIA window width in m/z ranges with less precursor ions present (12,14,15). MS2 fragment ions are measured subsequently to acquire high-resolution composite MS/MS spectra that are later deciphered by data processing and comparing DIA-MS data against spectral libraries, also referred to as reference libraries.

Multiple computational workflows dedicated to DIA-MS have been developed to accurately detect and quantify peptides and proteins from multiplexed DIA-MS/MS spectra (13, 14). Various strategies for data processing can be used: for example, the generation of sample specific, deep spectral libraries, that are generated by data-dependent acquisition (DDA) using the corresponding study samples on the same mass spectrometer as used for the quantitative DIA-MS, this experimental design is presented in this current study. Other strategies may involve the use of pre-existing spectral libraries (from other labs or previously generated), including large pan-human libraries (16), or even library-free approaches, where spectral libraries are directly built from DIA-MS files (15, 14). In general, these spectral libraries contain comprehensive peptide MS/MS information, such as precursor and fragment ions m/z, charge states, fragment ion intensities and distributions, chromatographic retention time and more (16). Acquired DIA-MS data is subsequently searched against the spectral libraries for peptide identification and quantification using mass spectrometric and chromatographic information to extract and score ion chromatograms of corresponding peak groups using a variety of target-decoy algorithms to determine false discovery rate (FDR) scores or q-values (12,16–18).

DIA-MS workflows are established on multiple different mass spectrometry (MS) platforms (19). In this study, we used a novel fast-acquisition rate SCIEX platform, an orthogonal quadrupole time-of-flight (QqTOF) platform, referred to as ZenoTOF 7600 system. Typically, QqTOF instruments exhibit very high acquisition rates in MS1 and MS/MS mode and are, therefore, particularly efficient for SWATH DIA workflows in highly complex matrices, such as organ tissues. However, traditionally, the orthogonal injection of ions from the quadrupole Q2 collision cell into the TOF flight tube typically suffers from an inefficient duty cycle, often with only 5–25% of ions transferred during each TOF pulse. A new technology, adding the Zeno trap, significantly improves the ion transmission at the TOF accelerator to ≥90% duty cycle across the mass range, by timing the ion-release from the Zeno trap with the pulse in the accelerator (**S1 Fig**) (20). The Zeno trap feature results in ions of all sizes arriving in the center of the TOF accelerator at the exact same time, where they are then efficiently pulsed into the TOF. This greatly improves sensitivity (with increases in sensitivity of 5–10-fold, depending on fragment ion m/z) with no reported loss in acquisition speed or spectral resolution. Markus Ralser’s group recently published an analysis of a K562 human cell lysate using this Zeno trap technology to acquire DIA-MS data (also termed Zeno SWATH DIA), in combination with fast chromatographic gradients, which highlighted the speed and sensitivity of the ZenoTOF 7600 system (21).

In this study, we used the novel ZenoTOF 7600 system, to assess its performance and DIA-MS capabilities for <u>quantification in a tissue organ (kidney)</u>, which poses specific challenges, due to the larger dynamic range of proteins in tissue matrices. We investigated the changes in the kidney proteome after AKI induced by ischemia-reperfusion in a high-throughput manner. We evaluate proteomic data collected from injured and contralateral control kidneys, applying three microflow chromatography gradient lengths from 20- to 120-min to gain insights into the relations between high-sample throughput, peptide identification efficiency, and quantification accuracy of significantly altered proteins in the injured kidney lysates. Finally, we present kidney proteome alterations in injured and control kidney to highlight affected biological pathways, including senescence-associated secretory phenotype markers that suggest an increased senescence burden upon AKI. In addition, we link these molecular changes with phenotypic mouse behavior changes and physiological measurements after AKI. Understanding proteome-wide differences induced by AKI will allow scientists to develop novel and more effective therapeutics for AKI.

## Results

### Acute Kidney Injury Model and Morphology

Four mixed background male mice (129/Black 6) were subjected to a renal ischemia-reperfusion injury by unilateral clamping of the left kidney pedicle for 26 min. In the model, one kidney was injured while the contralateral kidney was uninjured to serve as a healthy control. On Day 6 post-injury, the contralateral kidney was removed and processed for analysis. On Day 7, the mice were euthanized to harvest the injured kidney and blood was collected for analysis of kidney function. After harvesting, the kidney tissues were stained with hematoxylin and eosin (H&E) to assess morphology changes after renal damage (**Fig 1**). Kidney tubules of the contralateral control kidney tissue showed uniform morphology, intact tubules, and distinct nuclei. Cells in the injured kidney revealed disordered tubules with disrupted cell walls, sloughing of cellular contents, proteinaceous casts and damaged nuclei. H&E staining visually confirmed that the renal ischemia-reperfusion injury model selectively damaged one kidney while preserving the other as healthy tissue.

**Fig 1.**
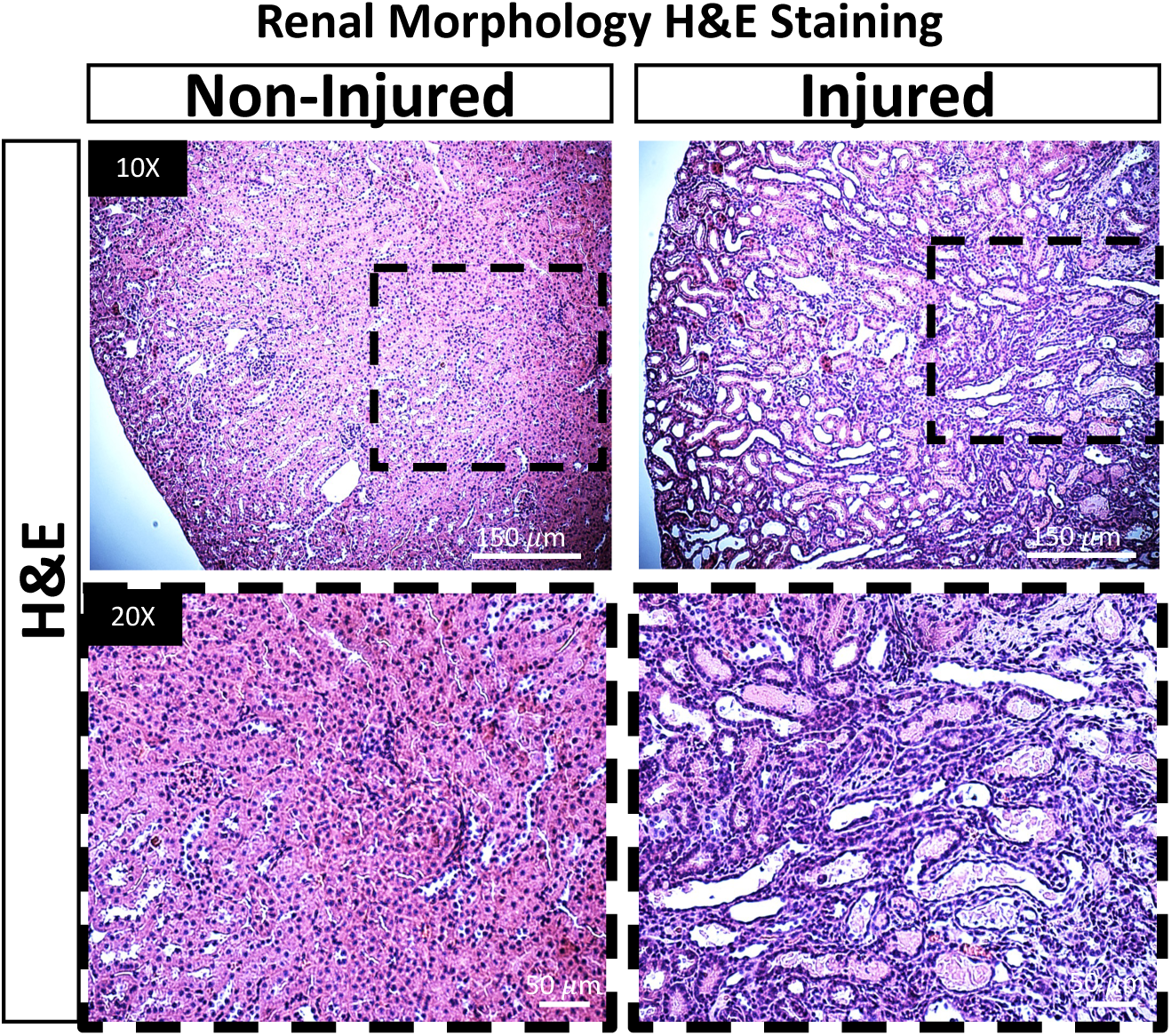
Ischemia-reperfusion promotes renal damage. Renal morphology of non-injured (control) and injured kidneys as shown by hematoxylin and eosin (H&E) staining. The control kidney tissue (panels on the right) is characterized by distinct and robust renal tubules, uniform cellular distribution, and intact cell walls. The injured kidney tissue (panels on the right) features damaged tubules and aggregated nuclei, non-uniform cellular distribution, disrupted cell walls, sloughing of cellular contents and proteinaceous casts. The panels on the top show a 10X zoom and the bottom panels show a 20X zoom in images of the indicated areas in the upper panels.

### Assessing Injured Kidney Injury Protein Markers

Creatinine and blood urea nitrogen (BUN) levels are classical clinical markers of kidney function: elevated levels indicate injury. Serum was collected at time of euthanasia and assessed by the Kansas State Veterinary Diagnostic Laboratory for creatinine and blood urea nitrogen (BUN) levels, compared to sham-surgery control animals (**Fig 2A-B**). In fact, both markers were highly elevated, serum creatinine was elevated by 10.44-fold (p<0.0001), and BUN was elevated by 7.95-fold (p<0.0001). In addition, using our novel mass spectrometric DIA-MS assay, we measured two additional kidney injury markers, kidney injury marker 1 (Kim-1) and neutrophil gelatinase-associated lipoprotein (NGAL), to confirm kidney injury (**Fig 2C-D**). Kim-1 was elevated by 7.13-fold (p=2.11 E-06), and NGAL was elevated by 4.76-fold (p=5.55 E-06). As expected, these established markers of kidney injury were significantly greater in the injured mouse kidney than in the contralateral healthy kidney indicating a very strong injury phenotype in our mouse model.

**Fig 2.**
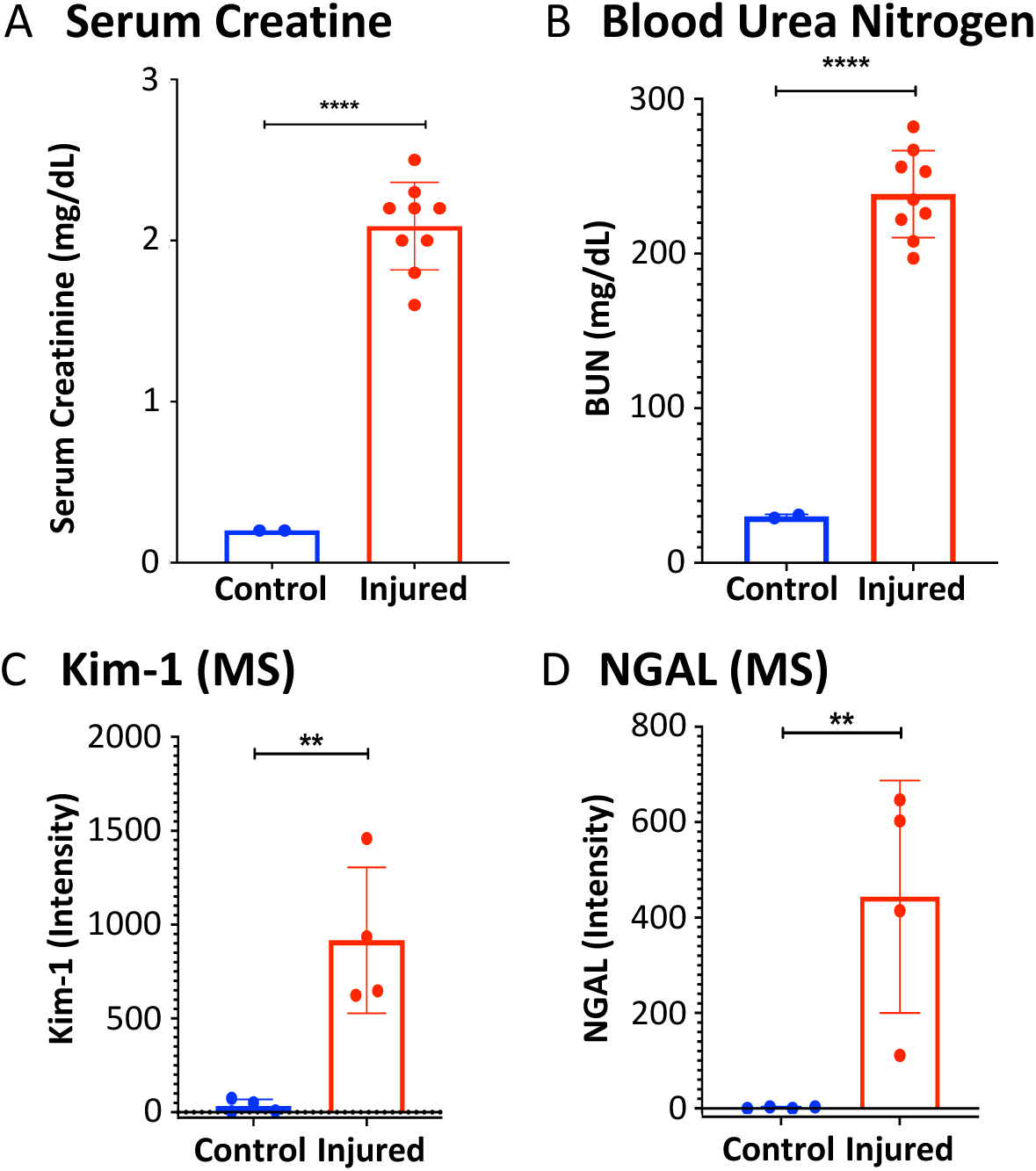
Injured kidney is elevated in markers of kidney injury. A) The serum biochemical levels of the kidney injury markers creatinine and B) blood urea nitrogen (BUN); and the MS2 intensity of the kidney injury markers C) kidney injury marker 1 (Kim-1) and D) neutrophil gelatinase-associated lipocalin (NGAL) are displayed. Results are expressed as mean +- S.D. Prism 9.0.0 software (GraphPad) was used for statistical analysis. Analysis was performed using Student’s t test. Significance was assigned by a p value < 0.05. *p<0.05 **p<0.01 ***p<0.001 ****p<0.0001.

### Proteomic Workflow and Generation of a Kidney-specific Deep Spectral Library

Flash-frozen kidney tissue was subjected to cell lysis and protein digestion with trypsin for mass spectrometric analysis using the ZenoTOF 7600 system (**Fig 3**). While some spectral libraries already existed for mouse kidney (22), we decided to build our own custom deep spectral library to be used for quantification of our future large-scale studies generating DIA-MS assays, monitoring AKI and future potential therapeutic interventions. Generating a spectral library unique to the kidney tissue from the mice used in this ischemia-reperfusion injury model with the ZenoTOF 7600 provides us confidence that the fragmentation will be consistent between the DDA and DIA data, and that the quantification accuracy of our assay is robust. Three technical replicates of each of the proteolytically digested injured (n=4) and contralateral healthy control (n=4) kidney samples with spiked-in iRT peptides were acquired with a ZenoTOF 7600 system using a Top 60 DDA workflow with Zeno trap activated using a 120-min microflow gradient to generate a deep spectral library in Spectronaut v16. Using a 120- min gradient length, each of the 4 injured and 4 contralateral control kidney lysates were acquired in technical triplicates in DDA mode, and the total of 24 acquisitions was used to build the new spectral library. This custom library that will be provided as a resource for the field contains 43,585 peptides and 4,062 protein groups with each containing two or more unique peptides per protein (**S1 Table**).

**Fig 3.**
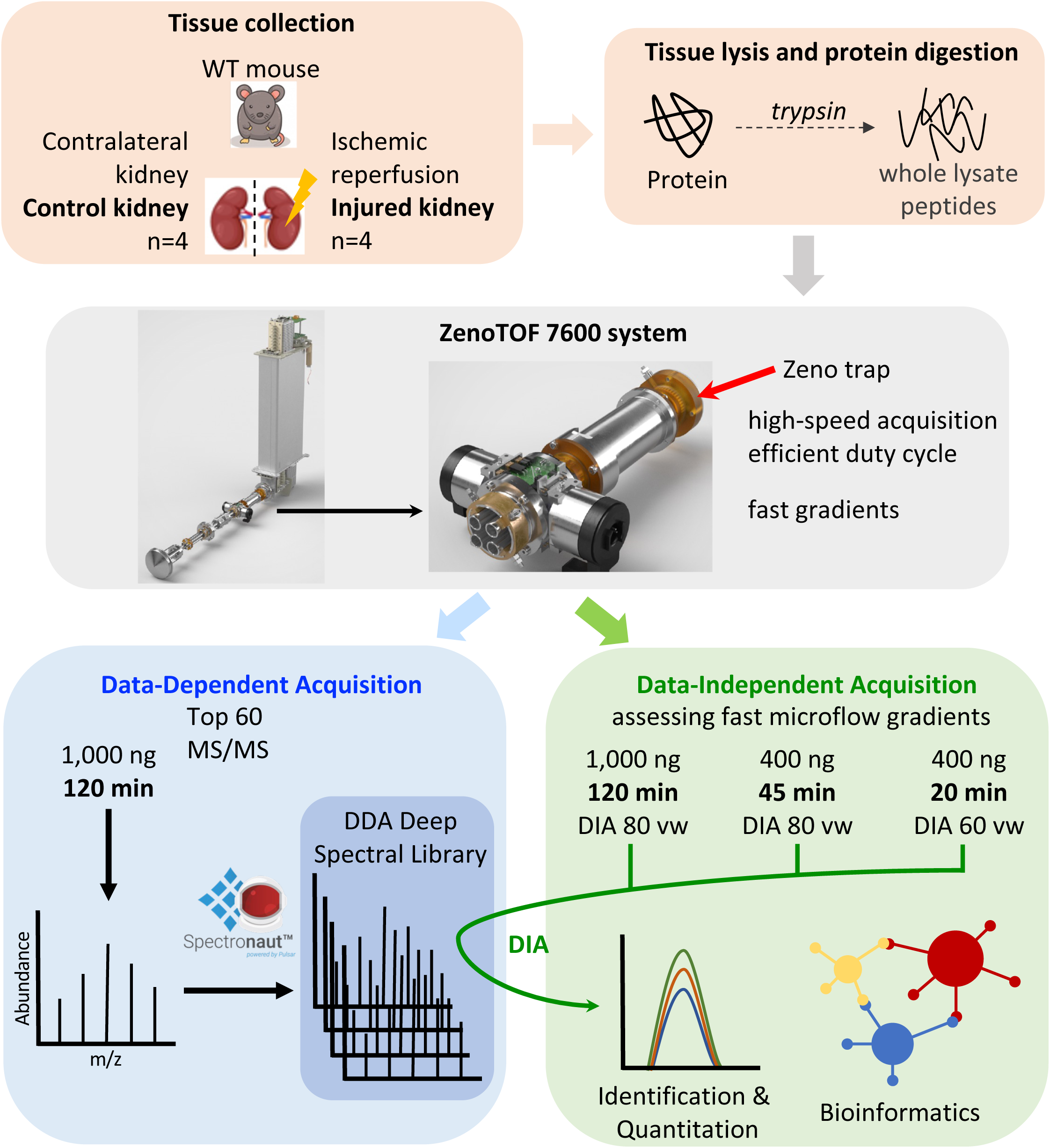
Main Proteomic Workflow. Ischemia-reperfusion AKI was performed on four wild-type (WT) mice for 22 min, and the contralateral kidney was preserved for use as a healthy control. The kidney tissue was homogenized, lysed, and prepared via tryptic digestion for mass spectrometric acquisition. The ZenoTOF 7600 system, which incorporates the novel Zeno trap technology to increase MS2 sensitivity, was coupled to an Acquity UPLC M-class system that allows for microflow chromatography and fast gradient separation of peptides. Data-dependent acquisition (DDA) of 1 µg of each kidney sample was performed in triplicate using a 120-min microflow gradient for the generation of a deep spectral library using Spectronaut v16 to generate a deep spectral library (referred to as ‘120-min library’). We acquired DIA-MS assays of each kidney sample injecting either i) 1 µg of material using a 120-min gradient with 80 DIA variable windows, ii) 400 ng of material using a 45-min gradient with 80 DIA variable windows, or iii) 400 ng of material using a 20-min fast gradient adjusting to 60 DIA variable windows. DIA-MS data were searched against the 120-min DDA deep spectral library for protein identification and quantification before bioinformatics analysis to compare the injured and control kidneys to identify protein candidates as potential biomarkers of AKI and to compare the efficiency of each microflow gradient.

### Proteome Changes Comparing Injured and Contralateral Healthy Kidney

Understanding the proteome changes comparing injured and healthy kidneys is essential for AKI biomarker discovery and development of treatments to mitigate AKI. A total of 3,945 protein groups were identified with two or more unique peptides in the DIA data using a chromatographic gradient of 120-min. Principal component analysis delineates a separation of the injured mouse kidneys and the contralateral healthy kidneys (**Fig 4A**). Differential protein quantification, comparing the proteolytically digested protein lysates from injured (n = 4) and control (n = 4) kidneys, demonstrated a complete remodeling of the kidney proteome upon acute injury. The volcano plot, shown in **Fig 4B**, highlights the statistically significant 1,285 up-regulated proteins in the injured mouse kidney and the 941 down-regulated proteins in the injured mouse kidney. The biological significance of these dramatic protein changes will be discussed below.

**Fig 4.**
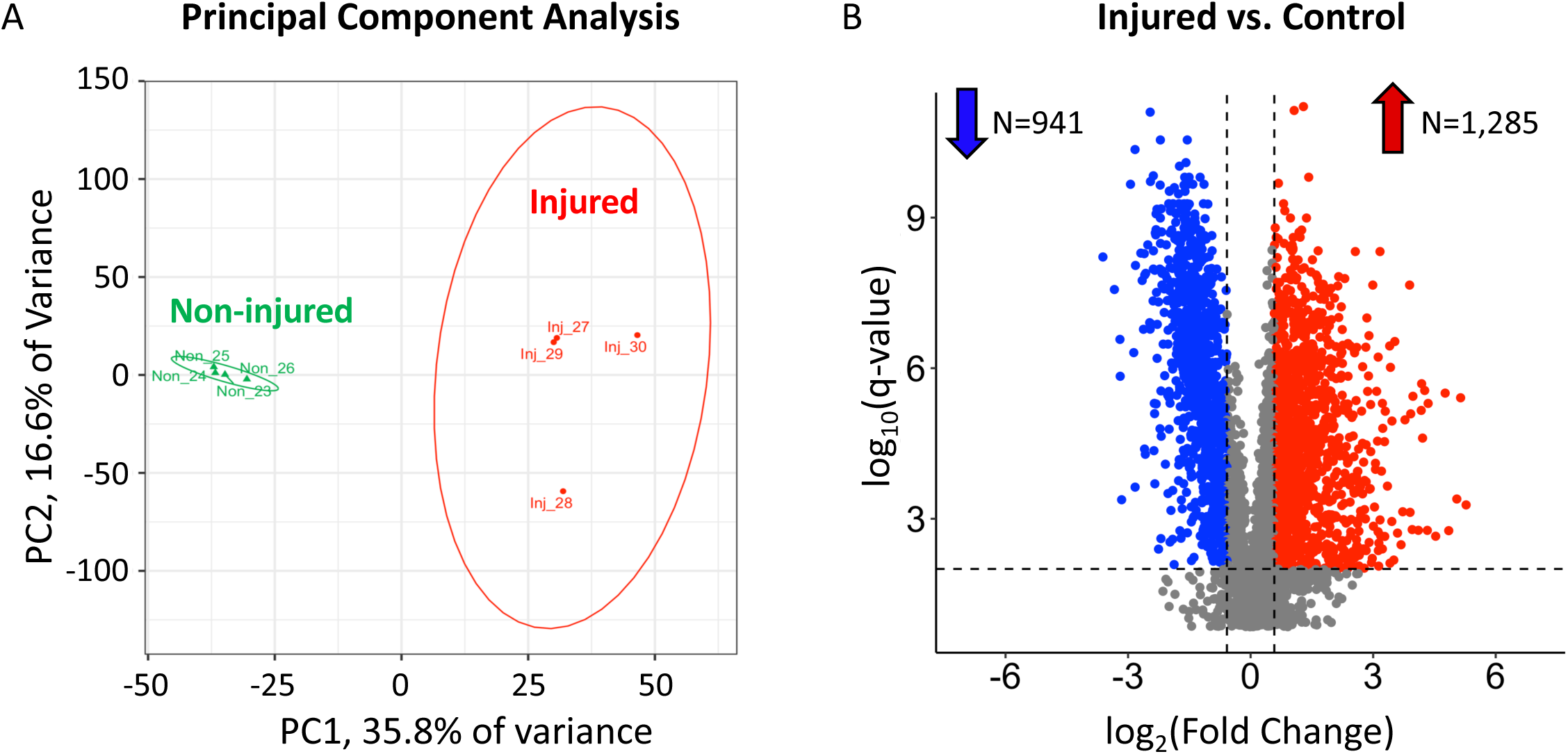
Ischemic-reperfusion completely remodels the kidney proteome. A) The principal component analysis of protein abundances for the injured and non-injured kidney indicates clear separation of the conditions. B) All 3,945 protein groups are displayed with significantly altered proteins (|log2(FC)| ≥ 0.58 & q-value < 0.01) highlighted in a volcano plot for the DIA-MS data acquired on the ZenoTOF 7600 system using a 120-min microflow gradient with a DIA isolation scheme of 80 variable windows. The 1,285 up-regulated proteins are shown in red, the 941 down-regulated proteins are shown in blue, and the 1,719 non-significantly altered proteins are shown in grey.

When comparing the proteome from the injured and contralateral control kidneys we observed a total of 941 down-regulated protein groups in the injured mouse kidney. Of these down-regulated proteins, the top 15 sorted by q-value are displayed in **S2A Table**. Of these 15 proteins, 11 proteins localize at least partially to mitochondria, consistent with reports that mitochondria are damaged by ischemic AKI (23). The top two proteins, peroxisomal acyl-CoA oxidase-2 (Acox2) and alpha-methylacyl-CoA racemase (Amacr), are key peroxisomal proteins involved in branched-chain fatty acid oxidation (FAO). Also downregulated was the trifunctional enzyme subunit alpha (HADHA), an important mitochondrial FAO enzyme. The top 15 down-regulated protein groups in the injured kidney, sorted by the log_2_ ratio of the fold-change, are displayed in **S2B Table.** These top-15 down-regulated protein groups have a significant function in key metabolic pathways, and their down-regulation after kidney injury drastically impacts kidney health.

When comparing the proteomes from injured (n = 4) and control (n = 4) kidneys we observed a total of 1,285 up-regulated protein groups. Of those the top 15 most statistically significantly up-regulated proteins are displayed in **S3A Table** (sorted by q-value). Ischemia-reperfusion of the kidney causes oxidative stress and DNA damage to the kidney proximal tubular epithelial cells. Proteins contributing to oxidative stress and DNA damage were more abundant in the injured kidney than the contralateral kidney. These proteins include Nucleolin (Ncl), prelamin-A/C (Lmna), palmitoyl-protein thioesterase 1 (Ppt1), microtubule-associated protein RP/EB family (Mapre1), serrate RNA effector molecule homolog (Srrt), and high-mobility group protein B1 (Hmgb1). Proteins involved in DNA repair or transcription have the highest significance (q-value ≤ 10^-9^) of the up-regulated proteins in the injured kidney and include Ncl (24), Lmna (25), Srrt (26), and Hmgb1 (27). Ppt1 is a hydrolase that participates in fatty-acid degradation in the lysosome, and the increased abundance indicates a shift in metabolism and energy production away from the mitochondria after kidney injury (28). The top 15 up-regulated protein groups in the injured kidney, sorted by the log_2_ ratio of the fold-change, are displayed in **S3B Table.** The up-regulated protein with the highest fold-change was hepatitis A virus cellular receptor 1 homolog (Havcr1), also known as kidney injury marker-1 (Kim1), a commonly used marker of kidney injury (29–32). Another marker of kidney injury, neutrophil gelatinase-associated lipocalin (NGAL or Lcn2), is also very high on the list of up-regulated proteins (33–36). The high abundance and upregulation of the kidney injury markers Kim1 and NGAL (7.13- and 4.76-fold, respectively) validated the use of mass spectrometry for the study of this renal ischemia-reperfusion model.

### Biological Processes affected Acute Kidney Injury

To perform biological pathway analysis of the 1,285 significantly up-regulated proteins and the 941 significantly down-regulated proteins that resulted from the ischemia-reperfusion in the injured kidney, we used ConsensusPathDB (37, 38). As ‘protein background’, we used all 4,062 identified protein groups that were also used to generate the deep spectral kidney library. The top 10 up- or down-regulated biological processes in injured kidney, when compared to the uninjured control kidney, are displayed in **Fig 5** (with q-value < 0.05 and term-level = 5). Up-regulated biological pathways in the injured kidney, including DNA conformation change, gene silencing by miRNA, positive regulation of response to DNA damage stimulus, and ATP-dependent chromatin remodeling, are likely related to oxidative damage to the DNA after ischemia-reperfusion (39, 40). Other up-regulated biological processes in the injured kidney, including B-cell-mediated immunity and negative regulation of coagulation indicate an immune response to ischemia-reperfusion (41). The down-regulated biological processes in the injured kidney are a direct response to ischemic injury, which include transmembrane transport of molecules in both the mitochondrial membrane and cell membrane, multiple metabolic and catabolic processes of fatty acids, and ATP production (8, 9).

**Fig 5.**
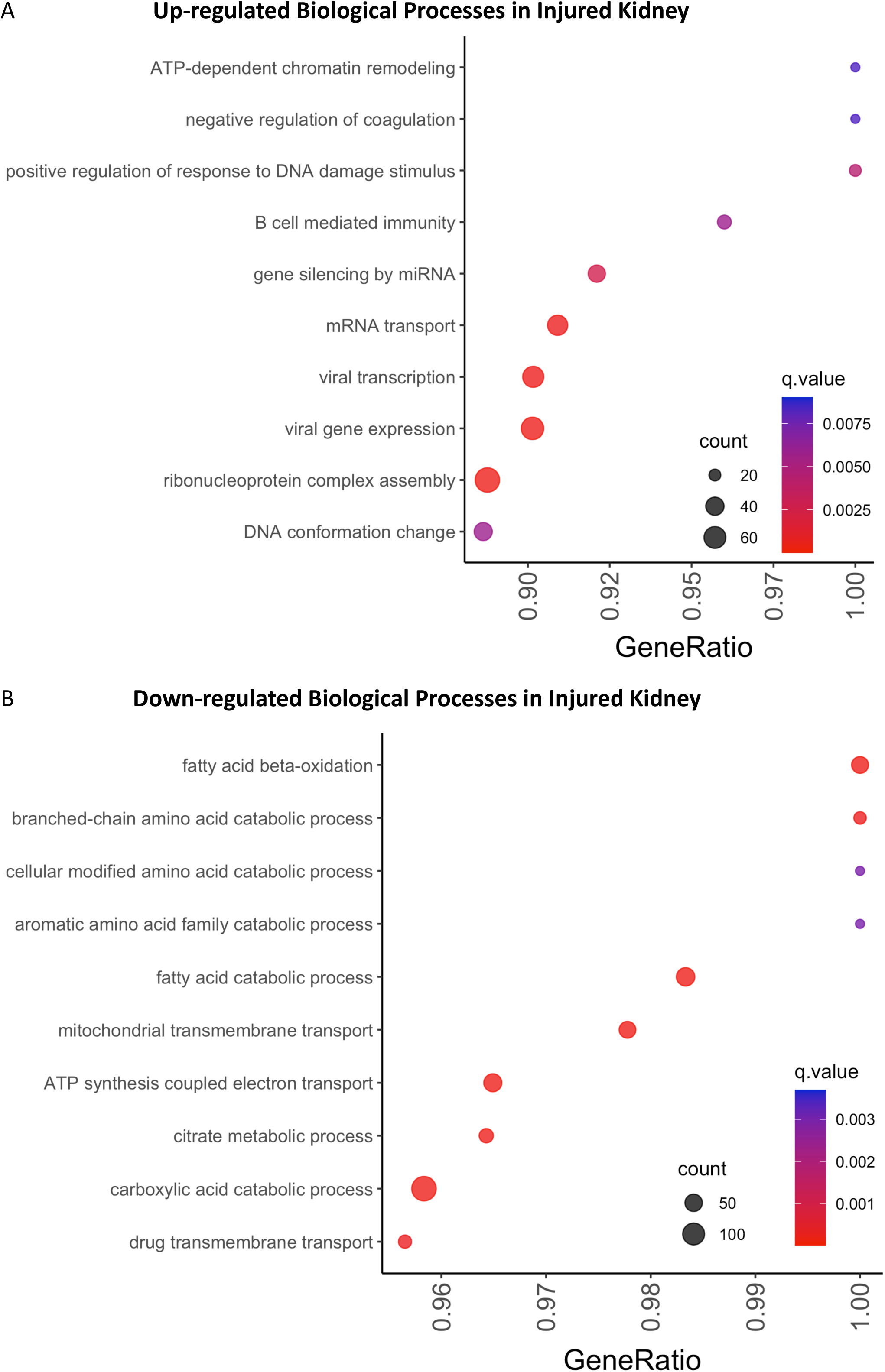
Metabolic responses are regulated by ischemia-reperfusion in the injured kidney. The top 10 biological processes are shown for the A) up- and B) down-regulated biological processes in injured kidney after searching the DIA data collected using a 120-min microflow gradient and 80 variable windows against the deep spectral library. The x-axis displays the ratio of the number of proteins identified as significantly altered in the biological process to the total number of proteins contributing to the biological process. The count is the number of significantly altered proteins contributing to the identification of the biological process. Biological processes were filtered to select for term category = b, a q-value < 0.01, and term level ≥ 5. The size of the dot indicates the number of protein groups that contribute to the identification of each biological process.

### Organelle Localization of Significantly Altered Proteins during Injured Kidney

To highlight organelles that are most impacted by AKI, we visualized the subcellular distribution of proteins using a methodology called “Localization of Organelle Proteins by Isotope Tagging” or LOPIT. The LOPIT methodology was originally developed by Kathryn Lilley’s group and combined biological fractionation by density-gradient ultracentrifugation, multiplexing with isobaric labels, and quantification by mass spectrometry to assign proteins to specific organelles (42). The discrete enrichment pattern of the proteomes of organelles that were separated into density-gradient fractions allowed assignment of proteins to their subcellular locations based on fraction and co-localization with proteins that define a specific organelle (43). A map of 5,032 proteins assigned to 15 subcellular locations in mouse pluripotent stem cell lysates from a LOPIT experiment conducted in the Lilley group were plotted using t-SNE coordinates as shown in **Fig 6A** (44, 45). Proteins that are present in multiple density-gradient fractions or that do not co-localize with protein markers of specific organelles occupy the central space between designated organellar locations, plotted in grey, on the t-SNE plot and have an undefined assignment. The injured vs. control kidney proteome fold-change data (**Fig 6B**) overlayed onto the LOPIT map demonstrate clear down-regulation of proteins (blue dots) in the mitochondria and peroxisome in the injured kidney after 26 min of ischemia-reperfusion and up-regulation of proteins (red dots) assigned to all other organelles in the injured kidney (44).

**Fig 6.**
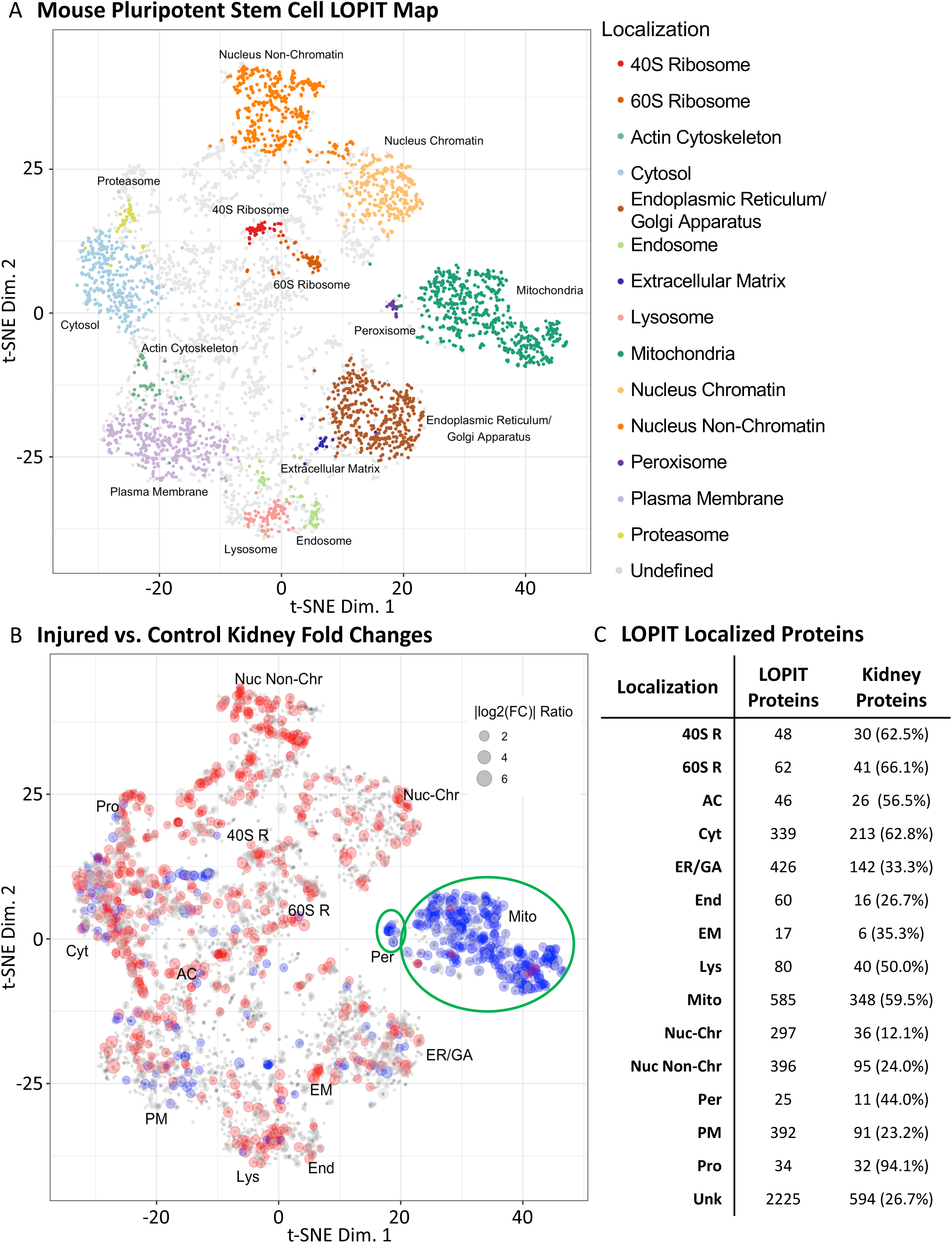
Organelle function is modified by ischemia-reperfusion in the injured kidney. A) The distribution of the LOPIT localized proteins for mouse pluripotent stem cells from the Lilley group is shown, *adapted from* Christoforou et al. 2016 (45). The assigned coordinates for 5,032 proteins in the 40S Ribosome (red, 40S R), 60S Ribosome (dark orange, 60S R), Actin Cytoskeleton (green, AC), Cytosol (light blue, Cyt), Endoplasmic Reticulum/Golgi Apparatus (brown, ER/GA), Endosome (light green, End), Extracellular Matrix (blue, EM), Lysosome (pink, Lys), Mitochondria (dark green, Mito), Nucleus – Chromatin (orange Nuc-Chr), Nucleus – Non-chromatin (light orange, Nuc Non-Chr), Peroxisome (purple, Per), Plasma Membrane (lavender, PM), Proteasome (yellow, Pro), and Undefined (grey) subcellular locations are shown. B) Binned injured vs. control log_2_ fold-change data are mapped to the LOPIT coordinates (1721 proteins) where upregulated proteins in the injured kidney are shown in red, and downregulated proteins the injured kidney are shown in blue. The dot size indicates the relative fold-change for each protein. Regions for mitochondria and peroxisome are highlighted with green ellipses. C) The total number of proteins localized to each organelle and the number of proteins quantified in the kidney lysates that are assigned to an organelle.

The LOPIT map of the fold-changes in injured kidneys shows how ischemia-reperfusion causes organelle-wide changes in the cells. The down-regulation of proteins localized to the mitochondria and peroxisomes is associated with disruptions in the metabolic processes necessary for energy production, In response, the cell may up-regulate proteins in the nucleus and ribosomes as a compensatory reaction. The increased production of nuclear proteins may be a response to cellular starvation, as the disruption of the Krebs cycle impairs the cell’s ability to produce energy. Overall, these changes highlight the complex effects of ischemia-reperfusion on the cellular metabolism and function.

### Fatty Acid Oxidation is Down-regulated in Injured Kidneys

As described above, the biological processes involved in fatty acid oxidation and metabolism are predominantly down-regulated in the injured kidney when compared to the uninjured control kidney. This is recapitulated in the LOPIT plots (**Fig 6B**), highlighting a strong observed down-regulation of protein groups in the mitochondria and peroxisome organelles. The numerous FAO proteins in the mitochondria and peroxisome pathways that are downregulated in the injured kidney are presented in **Fig 7** **and** **Fig 8**. **Fig 7** displays the fold-changes and q-values of the down-regulated mitochondrial proteins involved in catabolizing fatty acids, including carnitine palmitoyl- transferase-1 (CPT1A) (46, 47), carnitine palmitoyl-transferase-2 (CPT2) (48, 49), acyl-CoA synthetase (ACSM2 and ACSM3) (50), electron transfer flavoprotein (ETFA, ETFB, and ETFD) (51), acyl-CoA dehydrogenase (ACADL, ACADM, ACADS, ACADV, and ACDSB) (52), enolyl-CoA hydratase and reductase (DECR, ECH1, and ECHM) (53), 3- hydroxyacyl-CoA (ECHA) (54), 3-hydroxyacyl-CoA dehydrogenase (ECHA and HCDH) (55), and 3-ketocacyl-CoA thiolase (ECHB and THIM) (56). Sirtuin-3 and -5 are usually present in healthy kidney cells and help to maintain balance within the body (10). Sirtuins regulate various processes in the body, including oxidative stress, inflammation, and cell death, as well as the production of ATP and the development of mitochondria (10). However, in the injured kidney, the levels of these proteins are significantly down-regulated, which is consistent with observed decreases in the number of mitochondria and peroxisomes in the injured kidney.

**Fig 7.**
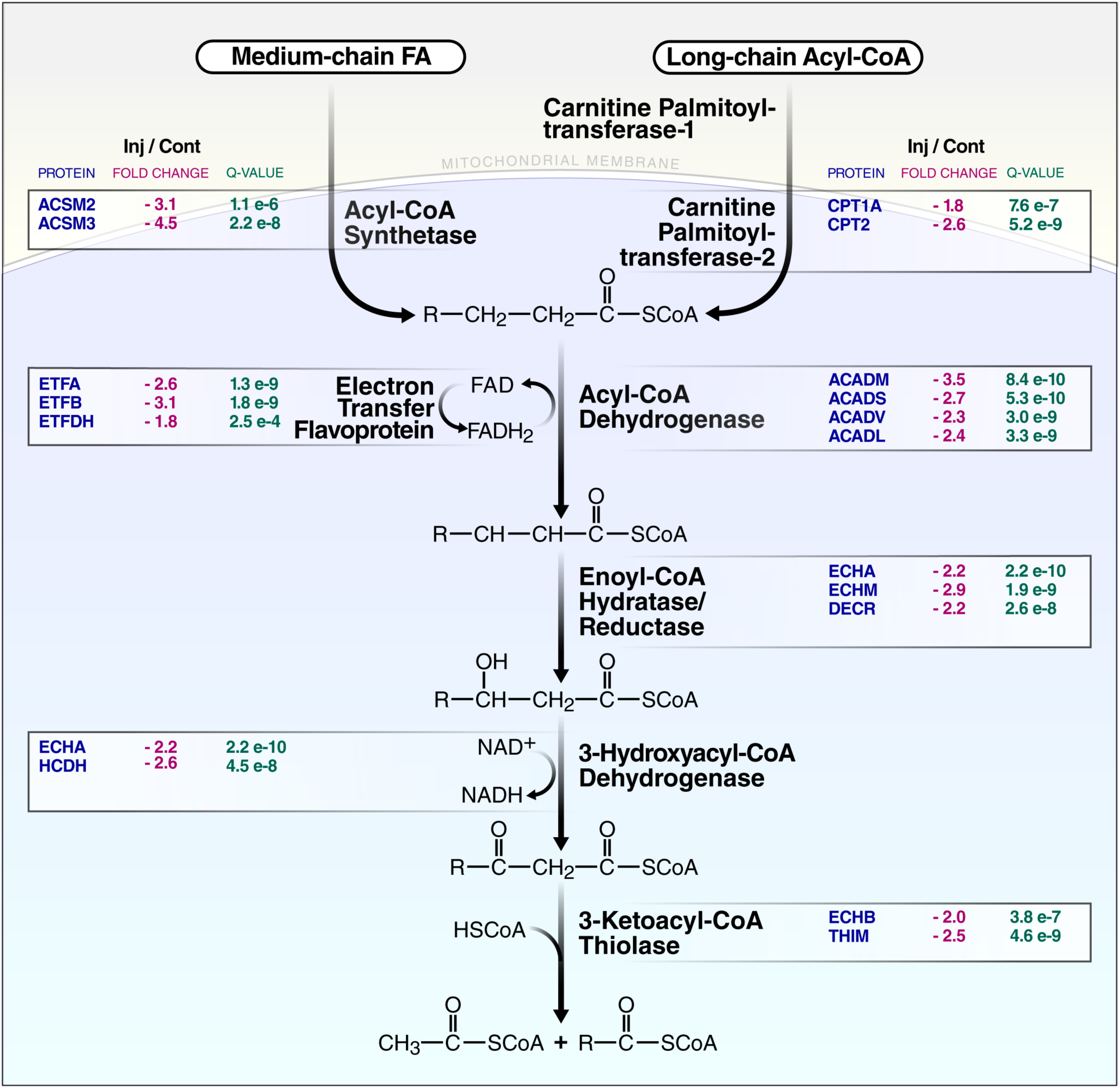
The fatty acid oxidation pathway is disrupted by kidney injury in the mitochondria. The proteins that were significantly down-regulated in the FAO pathway in mitochondria of the injured kidney are displayed with their fold-changes in blue and log_10_(q-values) in green (DIA-MS quantification employing the 120-min gradient).

**Fig 8.**
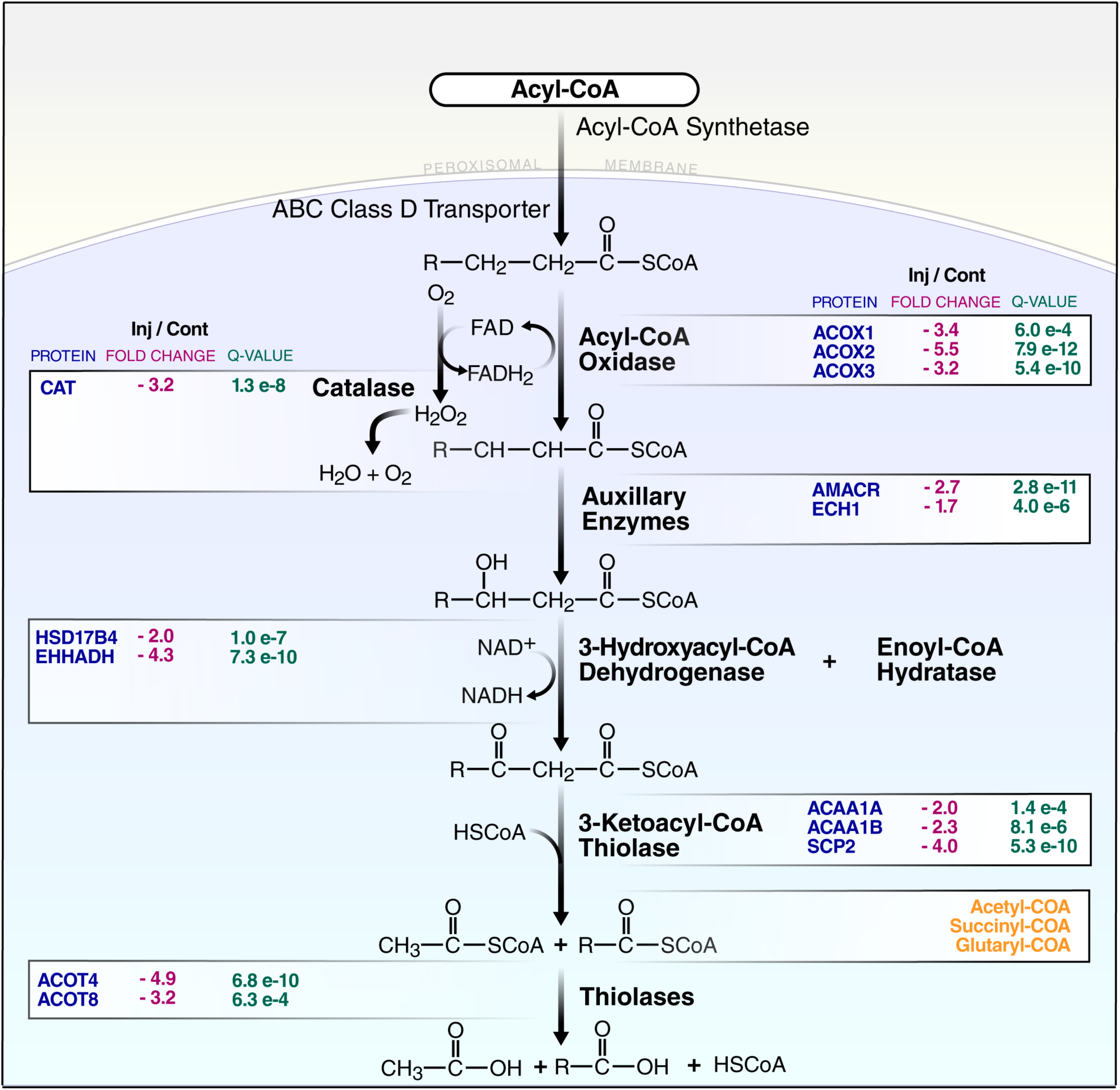
The fatty acid oxidation pathway is disrupted by kidney injury in the peroxisomes. Proteins that were significantly down-regulated in the FAO pathway in the peroxisome of the injured kidney are displayed with their fold-changes in blue and log_10_(q-values) in green (DIA-MS quantification employing the 120-min gradient).

Several peroxisomal proteins involved in catabolizing fatty acids are also down-regulated in the injured kidney (**Fig 8**). Down-regulated peroxisomal proteins include peroxisomal acyl-coenzyme A oxidase 1 (ACOX1) (57), catalase (CAT) (58), peroxisomal multifunctional enzyme type 2 (Hsd17B4) (59), peroxisomal bifunctional enzyme (EHHADH) (60), peroxisomal succinyl-coenzyme A thioesterase (ACOT4) (61), acyl-coenzyme A thioesterase 8 (ACOT8) (62), sterol carrier protein 2 (Scp2) (63), 3-ketoacyl-CoA thiolase A (Acca1a) and 3-ketoacyl-CoA thiolase B (Acca1b) (64). The PEX gene codes for a family of proteins that are involved in the transport of fatty acids across the membranes of cells, protein trafficking, peroxisome fusion and fission, and interactions between the peroxisome and endoplasmic reticulum (ER) (65, 66). These proteins are largely unaffected in injured kidney, suggesting that a selective loss of peroxisomal matrix proteins involved in metabolism occurs during AKI. The loss of specific peroxisomal proteins in the FAO pathway is extremely detrimental for the kidney as fatty acid molecules are still transported across the cell membrane and accumulate in the cytosol but are not processed in the mitochondria or peroxisome into the intermediates needed for energy production in the Krebs cycle.

### Many Senescence-Associated Secretory Phenotype Markers are Up-regulated after AKI

Cellular senescence, or permanent cell-cycle arrest, is a complex stress response that increases during aging and wound healing (67). Senescence occurs during the early stages of AKI and is a significant factor in the prognosis of the disease (68). Chronic senescence, which is characterized by changes at the transcriptional, metabolic, and secretory levels, as well as changes in cellular structure and chromatin organization, is particularly important in the progression of AKI to chronic kidney disease. Senescence can be improved through the process of reprogramming, which may prevent the progression of AKI to chronic kidney disease. Cellular senescence induces a complex senescence-associated secretory phenotype (SASP), including protein markers of inflammation, proteolysis, and extracellular matrix remodeling (69–72). The SASP markers are highly heterogenous depending on tissue type, but core SASP markers previously described by our group appear to indicate aging and disease mechanisms associated with permanent cell-cycle arrest (71). Almost half (49%) of all 151 core SASP markers were identified as significantly changed in injured kidney (**S2 Fig**).

To understand what biological processes are affected by the altered proteins that overlap with the Core SASP, the 67 up-regulated proteins in the injured kidney that overlap with core SASP markers were submitted to ConsensusPathDB over-representation analysis and searched with a background of all identified proteins (term level ≥ 4, q-value ≤ 0.01). The up-regulated biological processes of proteins that overlap with core SASP markers in the injured kidney include multiple processes related to extracellular matrix reorganization, such as cell junction assembly, wound healing, and morphogenesis of an epithelium. Several of the upregulated proteins in the injured kidney that are also SASP markers include collagens (Col12a1, Col6a1, and Col6a2) (73, 74), Serpin H1 (SerpinH1) (75), fibronectin (Fn1) (76), and periostin (Postn) (77). These extracellular matrix proteins are important in maintaining homeostasis in injured tissue and participate in wound repair and the overall healing process (78). Serpin H1, Fn1, and Postn are also up-regulated in chronic inflammation-associated cancer tissue, comparted to matched adjacent histologically normal tissue as described by Bons et al. (79). We speculate that their persistent up-regulation in the injured kidney may contribute to the progression from AKI to chronic kidney disease.

Of the significantly altered proteins that overlap with the core SASP, only seven were down-regulated in the injured kidney. Down-regulated core SASP markers in injured kidney include a mitochondrial 10-kDa heat shock protein (Hspe1) (80) and the enzymes lactoylglutathione lyase (Glo1) (58), aldo-keto reductase family 1 member A1 (Akr1a) (81), mitochondrial malate dehydrogenases (Mdh1 and Mdh2), superoxide dismutase(Sod1) (58), and fructose-bisphosphate aldolase C (Aldoc) (82). These down-regulated proteins are related to multiple metabolic processes and the tricarboxylic acid cycle. Overall, most significantly regulated proteins that also overlap with the core SASP were up-regulated in response to injury to bring the tissue back into homeostasis via wound repair, an important mechanism of the SASP.

### The Effect of Microflow Gradient Length for Protein Quantification and DIA-MS Assays

Typically, the sensitivity, depth and selectivity of the DIA-MS assays depend on the sample complexity, acquisition speed, and the DIA window isolation strategy. Subsequently, these factors affect the robustness of protein identifications and, more importantly, the accuracy of protein quantification. In addition, the MS1 precursor ion distribution of peptides generated from a tryptic digestion typically shows its highest density around mass-over-charge 500–700 m/z. Many modern DIA-MS acquisition methods employ variable MS2 window widths that take the precursor ion density distribution into account by applying smaller m/z windows in ‘busy’ m/z ranges (500-700 m/z) and larger m/z windows in ‘less busy’ m/z ranges as shown in **Fig 9A**, leading to less complexity, less interferences and more accurate quantification overall (15, 83). We acquired each of the digested injured (n = 4) and contralateral healthy control (n = 4) kidney samples using a ZenoTOF 7600 system, using a data-independent acquisition methodology (DIA) with Zeno trap activated, applying a 120-, 45-, or 20-min microflow gradient at 5 µL/min, respectively (**S3 Fig**). The DIA-MS data were acquired using a 80 variable windows isolation scheme for proteolytic mixtures separated using a 120- or 45-min microflow gradient. For the shorter 20-min microflow gradient a 60 variable windows isolation scheme was implemented. DIA-MS files for each gradient length were searched using the described custom deep spectral library in Spectronaut v16 to identify and quantify peptides and protein groups, and to generate pairwise comparisons between the injured and contralateral healthy control kidney samples (**S4 Fig, S5 Fig**).

**Fig 9.**
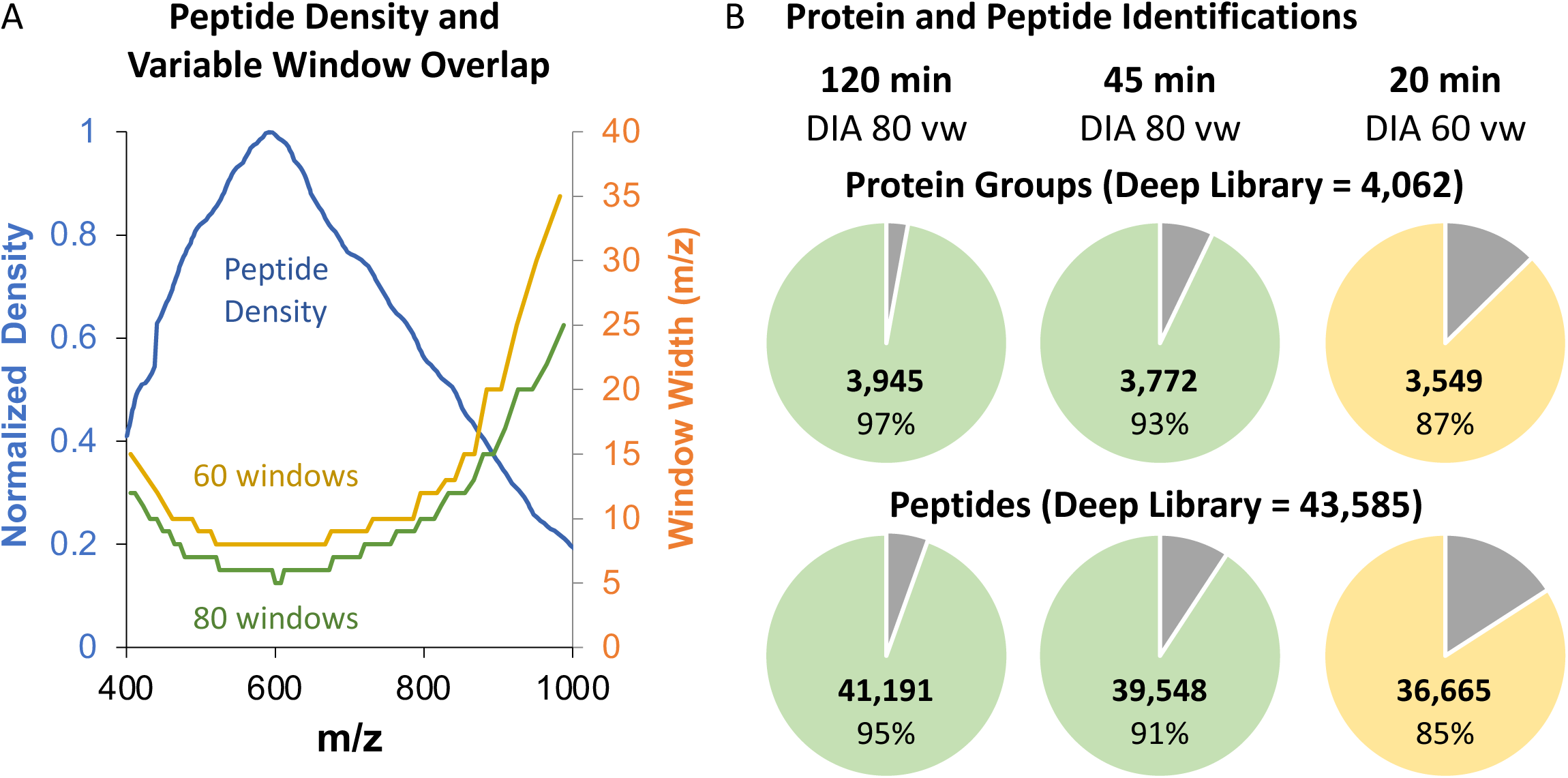
Efficient DIA-MS protein group and peptide identifications even when decreasing the length of the chromatographic gradient. A) The MS1 precursor ion density (blue) vs. m/z ratio is shown for tryptic digestions: for 60 variable windows (yellow) and 80 (green) variable windows are shown spanning a 400–1000 m/z range, that serves as m/z range during MS1 acquisition of the digested kidney tissue lysates. B) Pie charts display the number of protein and peptide identifications (color corresponds to variable windows) from the Zeno SWATH DIA data acquired using 20-, 45-, and 120-min gradients, respectively (yellow color indicates 60 variable windows and green color indicates 80 variable windows). The percentage in each pie chart represents the number of proteins and peptides quantified in the data relative to the total number identified in the custom deep spectral library. All data were searched with the deep spectral library, previously generated from DDA acquisitions using the 120-min gradient (grey).

All DIA-MS data of the injured (n = 4) and control (n = 4) kidney samples, acquired using a 120-, 45-, or 20-min chromatographic gradient, were searched against the deep spectral library described above, to detect and quantify peptides and proteins (**Fig 9B**). The Zeno SWATH DIA data (**S4 Table**) collected with the 120-min gradient (and 80 variable windows) resulted in the highest number of identifications and quantifications of 3,945 protein groups (97% of the total deep spectral library size) and 41,191 unique peptides (95% of the total deep spectral library size). Interestingly, the decrease of the gradient length from 120- to 45-min, using the identical DIA-MS method with 80 variable windows, showed similarly highly efficient identification and quantification performance, with 3,772 protein groups (93% of the total library size) and 39,548 unique peptides (91% of the total library size) quantified as shown in **S5 Table**. In addition, we were particularly interested in assessing even shorter gradients, and thus, we also acquired the Zeno SWATH DIA data of the proteolytically digested injured (n = 4) and contralateral healthy control (n = 4) kidneys using a relatively short 20-min gradient and a 60 variable window isolation scheme (**S6 Table**, searches used the identical above mentioned deep spectral library). Interestingly, these latter DIA-MS data resulted in similar numbers of identification and quantification of 3,549 protein groups (87% of the total spectral library size) and 36,665 peptides (85% of the total spectral library size). Thus, even short gradient lengths, combined with optimized Zeno SWATH DIA acquisition, proved highly efficient as 87% of all protein groups and 85% of all peptides were quantified using our deep custom kidney spectral library.

A comprehensive comparison of the Zeno SWATH DIA performance using either 120- or 20-min microflow gradients is presented in **Appendix 1** which highlights a strong similarity and correlation in data quality between the DIA data obtained using either a 120-min or 20-min microflow gradient. For instance, 3,500 protein groups with two or more unique peptides were quantified in both gradients. Also, the proteins that were identified as significantly altered in the DIA data collected with the 20-min gradient overlap with the proteins identified as altered in the DIA data obtained with the 120-min gradient. Additionally, the data obtained from the 20-min microflow gradient confirm the biological information previously presented for the DIA data collected with the 120-min microflow gradient. For example, the significantly altered proteins identified in the DIA data collected with a 20-min microflow gradient show a similar organellar distribution with down-regulation of proteins in the peroxisome and mitochondria. Submitting the significantly altered proteins from the DIA data obtained using a 20-min gradient to the ConsensusPathDB for gene ontology analysis resulted in the identification of similar up- and down-regulated biological processes. Up-regulated biological processes of the significantly up-regulated protein groups identified in the DIA data collected with a 20- min microflow gradient are involved in DNA damage, which is concordant with the up-regulated biological processes identified in the DIA data collected with the 120-min microflow gradient. Mitochondrial and peroxisomal biological processes of metabolism were identified as down-regulated in the injured kidney in DIA data collected with both the 20- and 120-min microflow gradients.

## Discussion

The rise in the incidence of patients with AKI is an important contributor to poor health outcomes that often can lead to chronic kidney disease and subsequent morbidity. The AKI mouse model presented in this work enabled the investigation of proteome differences between injured and contralateral healthy kidneys, while reducing biological heterogeneity and mouse to mouse variability through the preservation of one kidney as a contralateral healthy counterpart to the injured kidney. Orthogonal validation of kidney injury was achieved through a combination of histology, serum creatinine and blood urea nitrogen measurements, and mass spectrometry quantification of Kim-1 and NGAL. Our study provides novel insights into the molecular mechanisms of acute kidney injury and highlights distinct disease marker candidates, particularly those involved and dramatically decreased in fatty acid oxidation and energy metabolism.

We highlight that AKI completely remodels the kidney proteome, and over 50% of the protein groups quantified via DIA-MS showed significant changes in the injured kidney lysate, compared to the contralateral control. We observed a complete downregulation of fatty acid metabolism and energy production in the mitochondria and peroxisomes after insult by ischemic injury and reperfusion of the kidneys. The loss of mitochondrial function in the injured kidneys was observed via the downregulation of many key enzymes that are involved in the generation of acetyl-CoA. Loss of peroxisomal function in injured kidney was observed via the selective downregulation of peroxisomal matrix proteins involved in catabolizing fatty acids. These findings present exciting opportunities for the exploration of novel therapeutics to remediate mitochondrial and peroxisomal function after AKI.

The rise in AKI risk with increased patient age is well characterized in the literature (84, 85). Aging has been correlated to an increase in the burden of senescent cells (23), and we hypothesized that AKI may also cause an increase in the local tissue burden of senescent cells. Specifically, senescent cells and their SASP are involved in wound healing; however, the persistence of senescent cells may eventually also contribute to the transition from AKI to chronic kidney injury. In this study, injured kidneys were found to show elevated levels of over 49% of what we previously described as the ‘core SASP. Additional work remains to determine whether AKI increases the number of senescent cells persisting in kidney tissue after wound healing. However, our data offer the potential to use senolytics and senomorphics in the treatment of AKI to ameliorate senescent cell burden.

The novel ZenoTOF 7600 MS system incorporates the Zeno trap to significantly improve the duty cycle to nearly 100% for the orthogonal injection into the TOF region of the mass spectrometer, and to increase MS/MS sensitivity and overall quantification accuracy. Higher MS/MS sensitivity allows for much higher acquisition rates, and here, we demonstrated the coupling of accurate chromatographic microflow gradients with fast MS/MS acquisition, in data-dependent and data-independent workflows for efficient biological quantification studies. A deep kidney-specific spectral library, containing 43,585 peptides and 4,062 protein groups (>2 peptides), was generated using a ZenoTOF 7600 DDA method where 60 precursors were sent for MS/MS sampling in each cycle. This custom-built, deep spectral library has been made a publicly available resource to the scientific community, eliminating the time and cost of re-generating a spectral library while allowing for high-throughput and straightforward DIA assays for other kidney injury studies. The fast acquisition rates of the ZenoTOF 7600 platform enabled Zeno SWATH DIA methods using 80 or 60 variable windows to decrease spectral complexity and implement accurate protein quantification.

The ZenoTOF 7600 DIA-MS assays proved highly efficient to study AKI injury with accurate quantification, emphasizing interesting biology that changed upon kidney injury and that mostly impacted energy metabolism, including greatly reduced FAO performance in mitochondria and peroxisomes. However, we showed that, even when reducing the chromatographic gradient length from 120-to 45 or even 20-min, a very similar analytical performance highlighted the same remodeling of the kidney proteome upon injury and emphasizing the identical biological pathways affected. For future considerations, short gradient lengths may be especially needed and beneficial for larger cohort studies to reduce instrument time and increase sample throughput. The minimal loss in the number of peptides and protein groups quantified and the similar number of significantly altered protein groups quantified when reducing the gradient length from 120-min to 20-min indicate that the ZenoTOF 7600 system will serve as an excellent platform for large cohort studies or drug screening studies with desired high throughput.

Our study highlights the capabilities and potential of these novel high-throughput DIA-MS assay for assessing kidney injury in mice and other species. The MS assays presented above will allow for the monitoring of the proteome during drug development studies, including assessment of the dosage and time course in pre-clinical studies in a high-throughput manner. This is particularly useful for developing therapeutic interventions for acute kidney injury and can be extended to assays in humans. By monitoring changes in the kidney proteome as a response to injury and treatment, interventions may also be tailored to individual patients and their specific needs. Furthermore, the interventions developed for acute kidney injury could be extended to chronic kidney injury in human subjects. Our high-throughput MS assay allows for an efficient, comprehensive, and effective approach to studying and treating kidney disease.

## Materials and Methods

### Reagents

HPLC solvents (e.g., acetonitrile and water) were obtained from Burdick & Jackson (Muskegon, MI). Reagents for protein chemistry (e.g., triethylammonium bicarbonate buffer, iodoacetamide, dithiothreitol, and formic acid) were purchased from Sigma Aldrich (St. Louis, MO). Sequencing grade trypsin was from Promega (Madison WI). HLB Oasis SPE cartridges were purchased from Waters (Milford, MA). iRT peptides were purchased from Biognosys (Schlieren, Switzerland).

#### Ischemic AKI Models in Mice

B6/129SF1/J strain wild-type (WT) male mice were obtained from the Jackson Laboratory (stock no. 012757). Age matched 10–14-week-old mice were used throughout the study. The University of Pittsburgh Institutional Animal Care and Use Committee approved all experiments (approval no. 22112009). Mice (n = 4) were subjected to a renal ischemia-reperfusion injury model to induce ischemic AKI for 26 min as described, (86) with modifications. Briefly, mice were anesthetized with inhalant 2% isoflurane. The core body temperature was monitored throughout the procedures with a rectal thermometer probe and maintained at 36.8–37.2 °C with a heat lamp (Shat-R-Shield) and a water-heating circulation pump system (EZ-7150; BrainTree Scientific). Buprenorphine (Par Pharmaceutical) was administered subcutaneously for pain control (0.1 mg/kg body wt). A dorsal incision was made to expose the kidney and renal ischemia was induced for 26 min by unilateral clamping of the left kidney pedicle with a nontraumatic microvascular clamp (18055-04; Fine Science Tools) with aseptic techniques. Renal reperfusion was visually verified. The contralateral healthy control (n = 4) kidneys were harvested on Day 6. Mice were euthanized 7 days after injury to harvest the injured (n = 4) kidneys.

#### Morphology and Measurements of Kidney Injury Markers in Serum

For imaging: kidneys were fixed in 4% paraformaldehyde and embedded in paraffin before sectioning at 4 μm. The kidney sections were stained with hematoxylin and eosin (H&E) and subject to histologic evaluation, and ×10 and ×20 magnification images were obtained with a Leica DM2500 optical microscope. All tissue evaluations were performed in a blinded fashion. Blood from injured and sham-surgery control mice was collected by cardiac puncture, transferred to a tube containing a separator gel, and then centrifuged to collect serum. Serum analyzed by the Kansas State Veterinary Diagnostic Laboratory for creatinine and BUN levels.

#### Proteolytic Digestion

For mass spectrometric studies fresh frozen kidney tissue was used that were stored at −80 °C. For each individual kidney sample, 5.1 mg of protein lysate was suspended in 8 M urea in 50 mM triethylammonium bicarbonate buffer (TEAB, pH 8). Samples were reduced with 20-mM dithiothreitol in TEAB (pH 7) at 37 °C for 30 min, then cooled to room temperature (RT) and held at RT for 10 min. Samples were then alkylated with 40 mM iodoacetamide in 50 mM TEAB (pH 7) at RT in the dark for 30 min. The samples were diluted to 1 M urea using 50 mM TEAB (pH 7). Samples were incubated with sequencing grade trypsin (Promega, San Luis Obispo, CA) dissolved in 50 mM TEAB (pH 7) at a 1:50 (w/w) enzyme:protein ratio for 1 hour at 47 °C. Additional trypsin was added at the same w/w ratio, and proteins were digested at 37 °C overnight. Peptides were dried by centrifugal evaporation, resuspended in 0.2% formic acid (FA) in water and desalted with Oasis 30 mg Sorbent Cartridges (Waters, Milford, MA). The desalted elutions were dried via centrifugal evaporation and re-suspended in 0.2% FA in water at a final concentration of 1 µg/µL. Finally, indexed Retention Time Standards (iRT, Biognosys, Schlieren, Switzerland) were added to each sample, according to manufacturer’s instructions (87, 88).

##### Chromatographic Separation

All reverse-phase HPLC-MS/MS data were collected with a Waters M-Class HPLC (Waters, Massachusetts, MA) coupled online to a ZenoTOF 7600 system (SCIEX, Framingham, MA) with an OptiFlow Turbo V Ion Source (SCIEX) using a micro electrode (1–10 µL/min; SCIEX). The solvent system consisted of 0.1% FA in water (solvent A) and 99.9% acetonitrile (ACN), 0.1% FA in water (solvent B). Digested peptides were loaded onto a Luna Micro C_18_ trap column (20-x 0.30 mm, 5-µm particle size; Phenomenex, Torrance, CA) over 2 min at 10 µL/min with 100% solvent A. Peptides were eluted onto a Kinetex XB-C_18_ analytical column (150 x 0.30 mm, 2.6-µm particle size; Phenomenex) at 5 µL/min using a 120-, 45-, and 20-min microflow gradient (see below), respectively, each from 5 to 32% solvent B using the following paradigms (indicated the % of solvent B):

1. Digested peptides (1 µg) were loaded at 5% B, and separated using a **120-min linear gradient** from 5 to 32% B for 120-min, followed by an increase to 80% B for 1 min, a hold at 80% B for 2 min, a decrease to 5% B for 1 min, and a hold at 5% B for 6 min. The total HPLC acquisition length was 30 min. The total HPLC acquisition length was 130 min.
2. Digested peptides (400 ng) were loaded at 5% B, and separated using a **45-min linear gradient** from 5 to 32% B for 45 min, followed by an increase to 80% B for 1 min, a hold at 80% B for 2 min, a decrease to 5% B for 1 min, and a hold at 5% B for 6 min. The total HPLC acquisition length was 30 min. The total HPLC acquisition length was 55 min.
3. Digested peptides (400 ng) were loaded at 5% B, and separated using a **20-min linear gradient** from 5 to 32% B for 20-min, followed by an increase to 80% B for 1 min, a hold at 80% B for 2 min, a decrease to 5% B for 1 min, and a hold at 5% B for 6 min. The total HPLC acquisition length was 30 min.

##### Mass Spectrometric Data Acquisition and Processing

###### Data Acquisition

The source and gas parameters on the ZenoTOF 7600 system for all acquisitions were as follows: the ion source gas 1 was set to 10 psi, the ion source gas 2 was set to 25 psi, the curtain gas was set to 30 psi, the CAD gas was set to 7 psi, the source temperature was set to 200 °C, the HPLC column temperature was set to 30 °C, the polarity was set to positive, and the spray voltage was set to 5000 V.

###### Spectral Library Generation by Data-Dependent Acquisitions (DDA)

Three technical replicates each of the injured (n = 4) and control (n = 4) kidney lysate digests were collected with a DDA method using a 120-min microflow gradient with selection of the top, most abundant top 60 precursor ions per survey MS1 for MS/MS at an intensity threshold exceeding 100 cps (Top 60 DDA method). Sampled precursor ions were dynamically excluded for 10 s after one incidence of MS/MS sampling occurrence (MS/MS sampling with dynamic CE for MS/MS enabled). Precursor ions from charge states from 2 to 5 were selected for fragmentation using CID. The MS2 spectra were collected from 200–1,500 m/z with a 30-ms accumulation time and Zeno trap enabled. For the collision energy equation, the declustering potential was set to 80 V, with 0 V DP spread, and the CE spread was set to 0 V. The time bins to sum were set to 8 with all channels enabled and a 100,000 cps Zeno trap threshold. The cycle time for the Top 60 DDA method was 2.22 s.

###### Quantification by Data-Independent Acquisitions (DIA-MS)

For all gradient lengths, each of the samples were acquired in data-independent acquisition mode (Zeno SWATH DIA) as a single technical replicate with the Zeno trap enabled (11,12,15). The survey MS1 spectra were acquired from 395–1,005 m/z with a 100-ms accumulation time (**S3 Fig**). In the collision energy equation, the declustering potential was set to 80 V, with 0 V DP spread, the collision energy was set to 10 V with 0 V CE spread for efficient fragmentation of all precursor ions. The ‘time bins to sum’ were set to 8 with all channels enabled. In the DIA method using the 120- and 45-min microflow gradient, the MS1 accumulation time was set to 25-ms and 80 variable windows were used to collect MS/MS spectra, resulting in a total cycle time of 2.5-s. In the DIA MS method using the 20-min gradient, the MS1 accumulation time was set to 20-ms and 60 variable windows were used to collect MS2 spectra with a total cycle time of 1.617-s.

###### Spectronaut Mass Spectrometric Spectral Library Generation (DDA) with DIA-MS Data Processing, Quantification, and Statistical Analysis

All data files were processed in Spectronaut (version 16.0.220524.5300; Biognosys) using spectral library search for both the peptide and protein levels. Data were searched against the *Mus musculus* reference proteome with 58,430 entries (UniProtKB-TrEMBL), accessed on 01/31/2018. Dynamic data extraction parameters and precision iRT calibration with local non-linear regression were used. Trypsin/P was set as the digestion enzyme with specific cleavages and up to two missed cleavages allowed. Methionine oxidation and protein N-terminus acetylation were set as dynamic modifications, and carbamidomethylation of cysteine was set as a static modification. Identification at the protein group level required the at least two unique peptide identifications and was performed with a 1% q-value cutoff of the precursor ion and protein level. The protein level quantification was based on the peak areas of extracted ion chromatograms (XICs) of 3–6 MS2 fragment ions, specifically b- and y-ions with and automatic normalization strategy and 1% q-value data filtering applied. Relative protein abundance changes were compared in a statistically relevant manner using the Storey method with paired t-tests and p-values corrected for multiple testing by applying group wise testing corrections (89). **S1 Table** contains protein identifications from the custom deep spectral library. **S4 Table, S5 Table, and S6 Table** contain protein identifications and quantification of DIA data collected with a 120-, 45-, or 20-min microflow gradient, respectively.

###### Pathway Analysis

Consensus Path DB-mouse (Release MM11, 14.10.2021) was used for over-representation analysis (ORA) of the significantly altered quantifiable proteins (|log2(FC)| ≥ 0.58 & q-value < 0.01) to identify which gene ontology terms were significantly enriched in these samples (37, 38). Gene ontology terms (including biological processes, molecular functions, and cellular components) were filtered to select for biological processes (term category = b) with a q-value < 0.01 and term level ≥ 5. Dot plots were generated using the ggplot2 package (90) in R (version 4.0.5; RStudio, version 1.4.1106) to visualize significantly enriched biological processes from each comparison (**Fig 6**).

###### Localization of Organellar Proteins by Isotope Tagging (LOPIT) Map

Rstudio (v4.0.5, https://www.rstudio.com/ products/rstudio/download/#download, accessed on 31 March 2021) with R Bioconductor packages pRolocData and pRoloc were used to download the mouse pluripotent stem cell (hyperLOPIT2015) dataset from Christiforou et al. (2016)(91, 45). The hyperLOPIT2015 dataset contains biological organelle density fraction enrichment patterns, quantitative multiplexed MS data of each fraction, and protein localization assignments based on similarities in distribution to well-annotated organelle protein markers (43). The LOPIT map in **Fig 7** was created using the t-SNE machine learning algorithm to reduce the multi-dimensional hyperLOPIT2015 dataset to cluster proteins by similarities in the multiple experimental factors described above and overlaying the points with our own fold-change data comparing the injured and healthy kidney (44). All protein organelle assignments are used without additional refinement or alteration.

###### Programming Code Availability

The R scripts used in this study are available through our GitHub repository (https://github.com/JoBBurt/Kidney-Injury-ZenoTOF-DIA) and can be cited using the DOI: https://doi.org/10.5281/zenodo.7553648.

## Acknowledgements

This work was supported by the National Institutes of Health (NIH), specifically the National Institute of Diabetes and Digestive and Kidney Diseases (NIDDK) R01 DK090242 (to SSL) and R56 DK121758-01 (to ESG). The authors acknowledge the very generous support from SCIEX for the ZenoTOF 7600 system and a Waters M-class HPLC system at the Buck Institute.

## Supplementary Materials

**S1 Fig. The Zeno trap increases the efficiency of the duty cycle in the ZenoTOF 7600 system.** A) Zeno trap and TOF assembly. B) Zeno trap activated. C) Released based on potential energy such that large ions first and small ions last from the Zeno trap to the TOF accelerator. D) Pulsing all ions into the TOF at the same time.

**S2 Fig. Overlap in significantly altered proteins and the core SASP demonstrates upregulation of processes involved in wound healing.** This shows the overlap of proteins identified in the DIA data collected using an 80 variable window method and 120-min chromatographic gradient and the core senescence associated secretory phenotype (SASP) proteins. The core SASP proteins identified in the dataset are listed. Biological processes for up- and down-regulated SASP markers are shown in dot plots.

**S3 Fig. Zeno SWATH DIA data acquisition and processing settings.** Waters M-Class microflow chromatography hardware and flow rates; ZenoTOF 7600 system acquisition settings for DDA and DIA collected using a 120-, 45-, and 20-min microflow gradient; and plot of the overlap in peptide density and variable window coverage for 60 and 80 variable window methods.

**S4 Fig. Simplified proteomic workflow.** Scheme of the sample preparation, data collection, and data processing for the DDA data collected using a 120-min microflow gradient to build a custom deep spectral library and for the DIA data acquired using a 20-, 45-, or 120-min microflow gradient to identify peptides and proteins by searching against the deep spectral library for accurate quantification.

**S5 Fig. Quality assessment of DIA data collected with a 120-min microflow gradient.** Spectronaut 16 generated plots for retention time calibration, iRT peptide retention times, data completeness, and q-values of identifications demonstrate the robustness of the DIA data acquired with 80 variable windows using a 120-min microflow chromatography gradient.

**S1 Table. All proteins and peptides contained in the custom DDA deep spectral library.** Displaying all peptides and proteins identified in the custom DDA deep spectral library. DDA data was collected using the top 60 precursor selection per MS cycle using a 120-min microflow chromatography gradient.

**S2 Table. Up-regulated proteins in injured kidney from the DIA data collected with the 120-min microflow gradient.** Displaying the top 15 upregulated proteins from the DIA data collected with 80 variable windows using the 120-min microflow gradient, sorted by log_2_(fold-change), in the injured kidney.

**S3 Table. Down-regulated proteins in injured kidney from the DIA data collected with the 120-min microflow gradient.** Displaying the top 15 downregulated proteins from the DIA data acquired using the 120-min microflow gradient with 80 variable windows, sorted by log_2_(fold-change), in the injured kidney.

**S4 Table. Proteins identified in the DIA data collected with a 120-min microflow gradient.** Data for the 3,945 protein groups identified with 2 or more unique peptides and for the 2,226 significantly altered protein groups with a |log_2_(FC)| ≥ 0.58 and q-value ≤ 0.01 in the DIA data collected with a 120-min microflow gradient using 80 variable windows is displayed.

**S5 Table. Proteins identified in the DIA data collected with a 45-min microflow gradient.** Data for the 3,772 protein groups identified with ≥ 2 unique peptides and for the 1,867 significantly altered protein groups with a |log_2_(FC)| ≥ 0.58 and q-value ≤ 0.01 in DIA data collected with 80 variable windows using a 45-min microflow gradient is displayed.

**S6 Table. Proteins identified in the DIA data collected with a 20-min microflow gradient.** Data for the 3,549 protein groups identified with ≥ 2 unique peptides and for the 1,658 significantly altered protein groups with a |log_2_(FC)| ≥ 0.58 and q-value ≤ 0.01 in the DIA data collected with a 20-min microflow gradient using 60 variable windows is displayed.

