## Supplementary material for "Substantial Downregulation of Mitochondrial and Peroxisomal Proteins during Acute Kidney Injury revealed by Data-Independent Acquisition Proteomics": Suppl Figures: Figure_S-01_Zeno_Kidney_23_0224_v04.pptx

### Slide 1
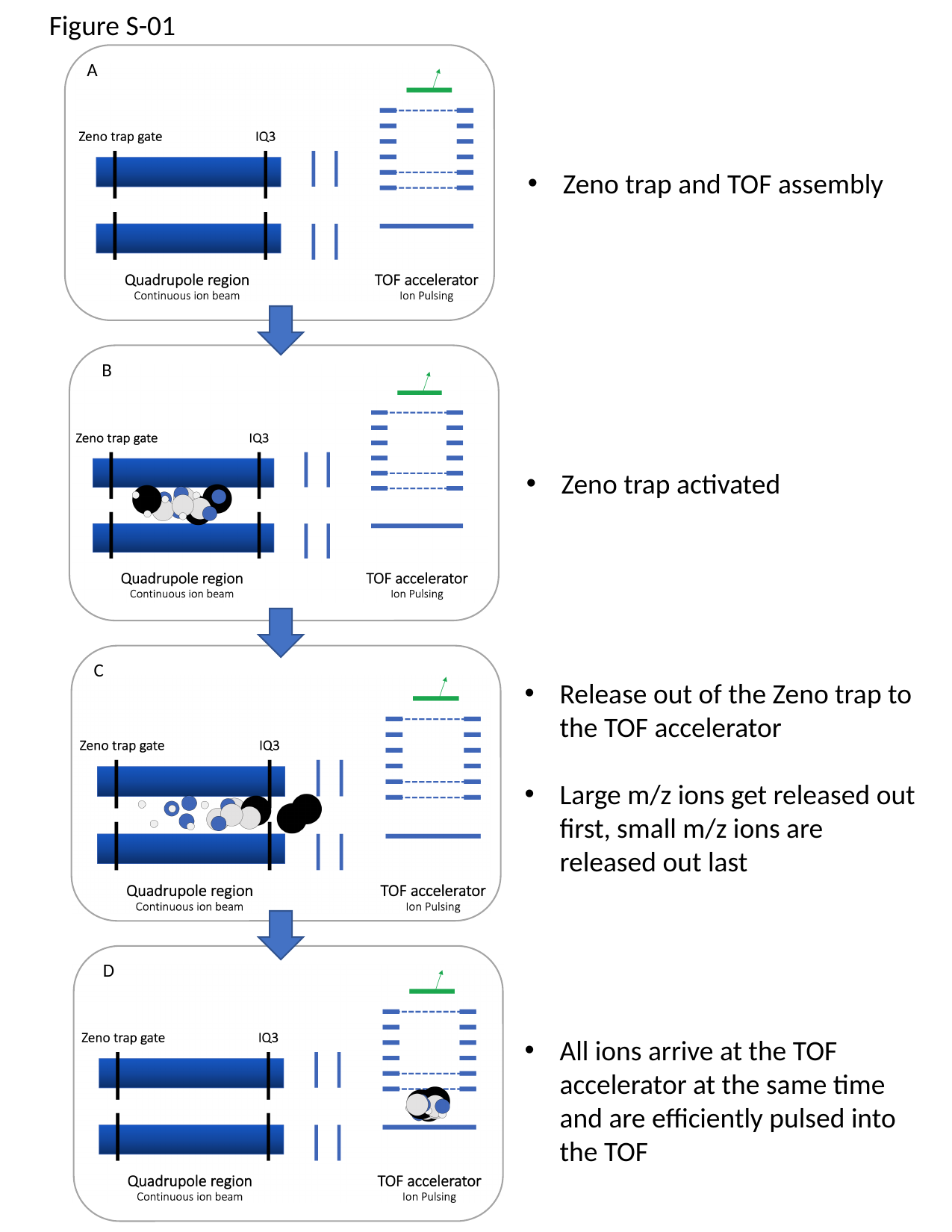

Figure S-01
A
Zeno trap and TOF assembly
B
Zeno trap activated
C
Release out of the Zeno trap to the TOF accelerator
Large m/z ions get released out first, small m/z ions are released out last
D
All ions arrive at the TOF accelerator at the same time and are efficiently pulsed into the TOF

### Slide 2
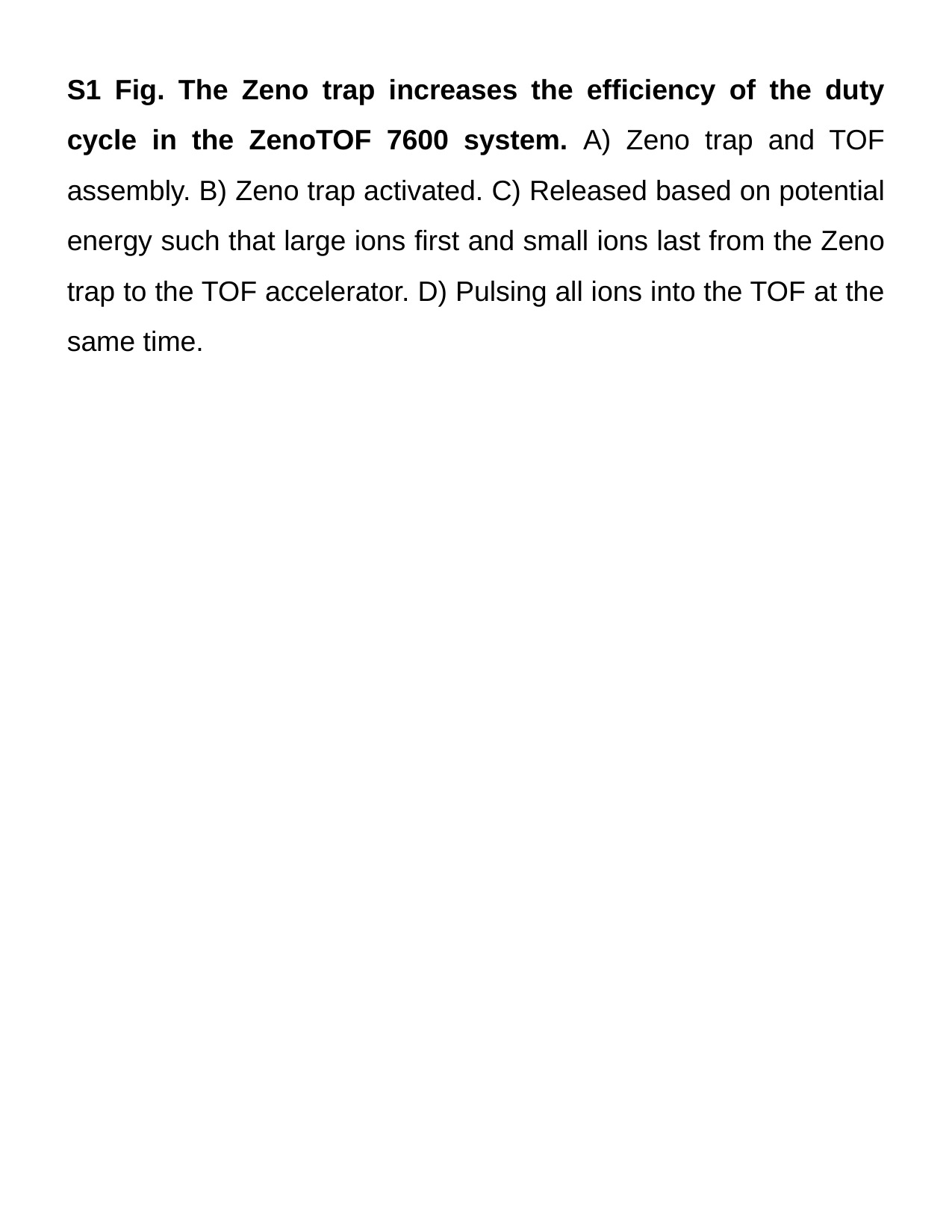

S1 Fig. The Zeno trap increases the efficiency of the duty cycle in the ZenoTOF 7600 system. A) Zeno trap and TOF assembly. B) Zeno trap activated. C) Released based on potential energy such that large ions first and small ions last from the Zeno trap to the TOF accelerator. D) Pulsing all ions into the TOF at the same time.
