## Supplementary material for "Substantial Downregulation of Mitochondrial and Peroxisomal Proteins during Acute Kidney Injury revealed by Data-Independent Acquisition Proteomics": Suppl Figures: Figure_S-02_Zeno_Kidney_23_0224_v05.pptx

### Slide 1
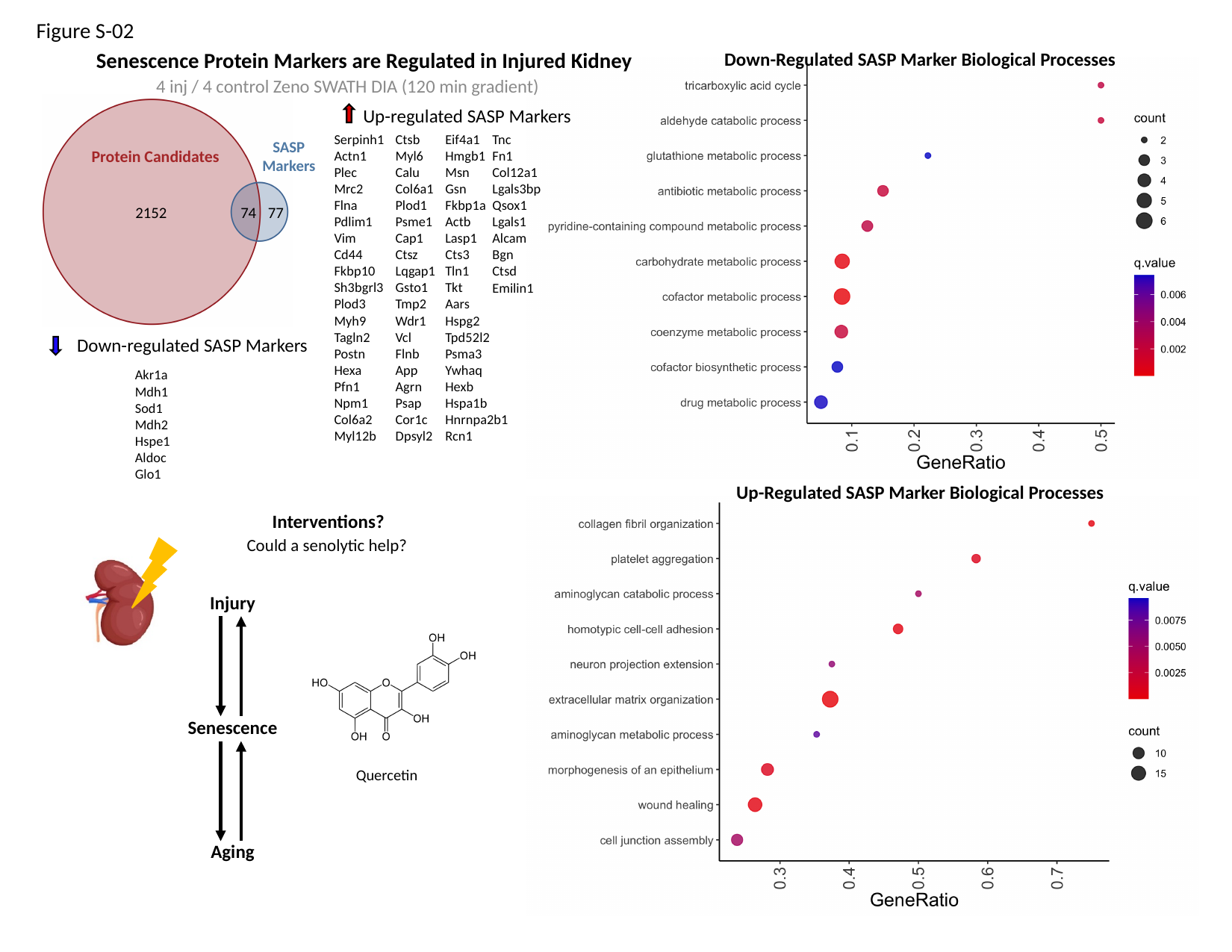

Figure S-02
Down-Regulated SASP Marker Biological Processes
Senescence Protein Markers are Regulated in Injured Kidney
4 inj / 4 control Zeno SWATH DIA (120 min gradient)
SASP
Markers
Protein Candidates
2152
74
77
Up-regulated SASP Markers
Serpinh1
Actn1
Plec
Mrc2
Flna
Pdlim1
Vim
Cd44
Fkbp10
Sh3bgrl3
Plod3
Myh9
Tagln2
Postn
Hexa
Pfn1
Npm1
Col6a2
Myl12b
Ctsb
Myl6
Calu
Col6a1
Plod1
Psme1
Cap1
Ctsz
Lqgap1
Gsto1
Tmp2
Wdr1
Vcl
Flnb
App
Agrn
Psap
Cor1c
Dpsyl2
Eif4a1
Hmgb1
Msn
Gsn
Fkbp1a
Actb
Lasp1
Cts3
Tln1
Tkt
Aars
Hspg2
Tpd52l2
Psma3
Ywhaq
Hexb
Hspa1b
Hnrnpa2b1
Rcn1
Tnc
Fn1
Col12a1
Lgals3bp
Qsox1
Lgals1
Alcam
Bgn
Ctsd
Emilin1
Down-regulated SASP Markers
Akr1a
Mdh1
Sod1
Mdh2
Hspe1
Aldoc
Glo1
Up-Regulated SASP Marker Biological Processes
Interventions?
Could a senolytic help?
Injury
Senescence
Quercetin
Aging

### Slide 2
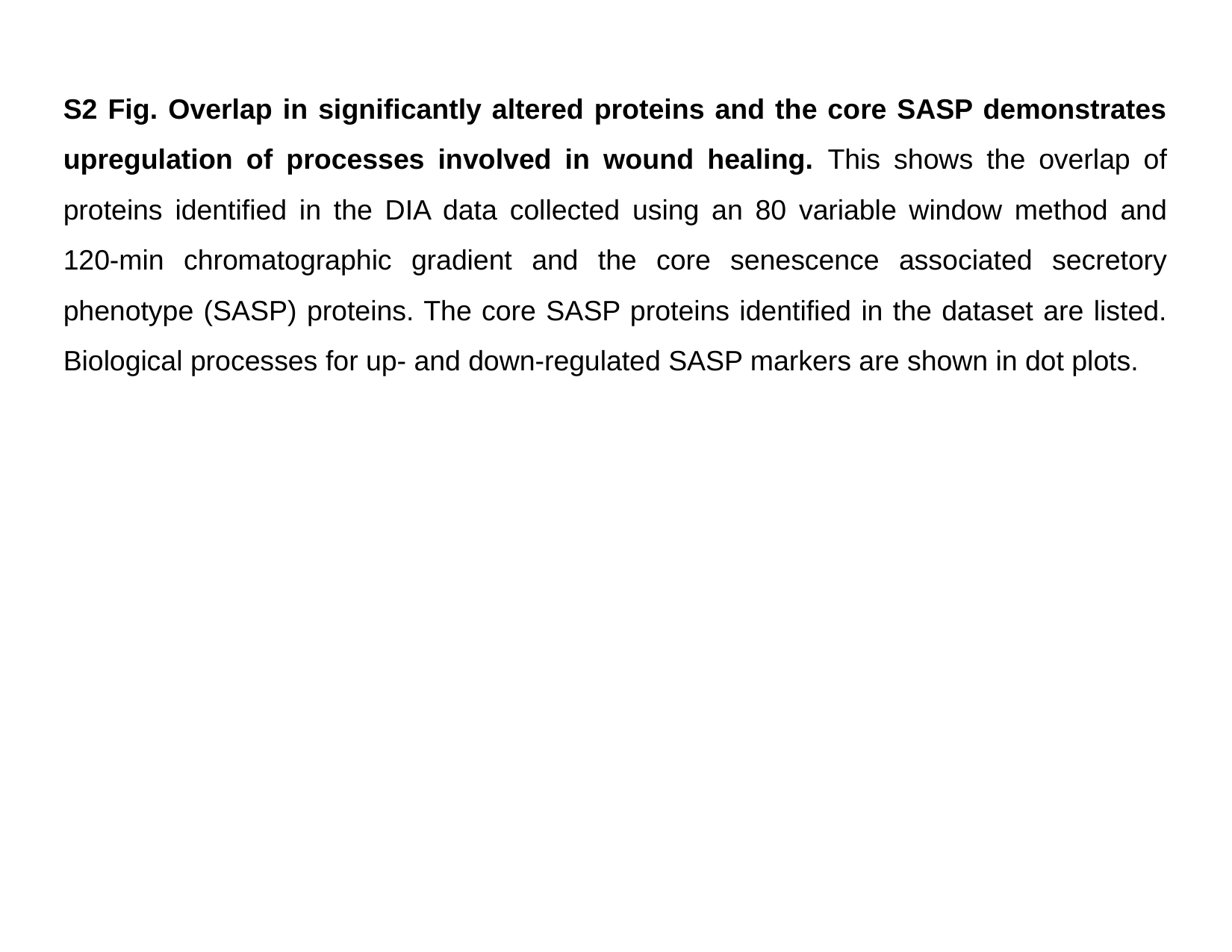

S2 Fig. Overlap in significantly altered proteins and the core SASP demonstrates upregulation of processes involved in wound healing. This shows the overlap of proteins identified in the DIA data collected using an 80 variable window method and 120-min chromatographic gradient and the core senescence associated secretory phenotype (SASP) proteins. The core SASP proteins identified in the dataset are listed. Biological processes for up- and down-regulated SASP markers are shown in dot plots.
