## Supplementary material for "Substantial Downregulation of Mitochondrial and Peroxisomal Proteins during Acute Kidney Injury revealed by Data-Independent Acquisition Proteomics": Suppl Figures: Figure_S-03_Zeno_Kidney_23_0224_v03.pptx

### Slide 1
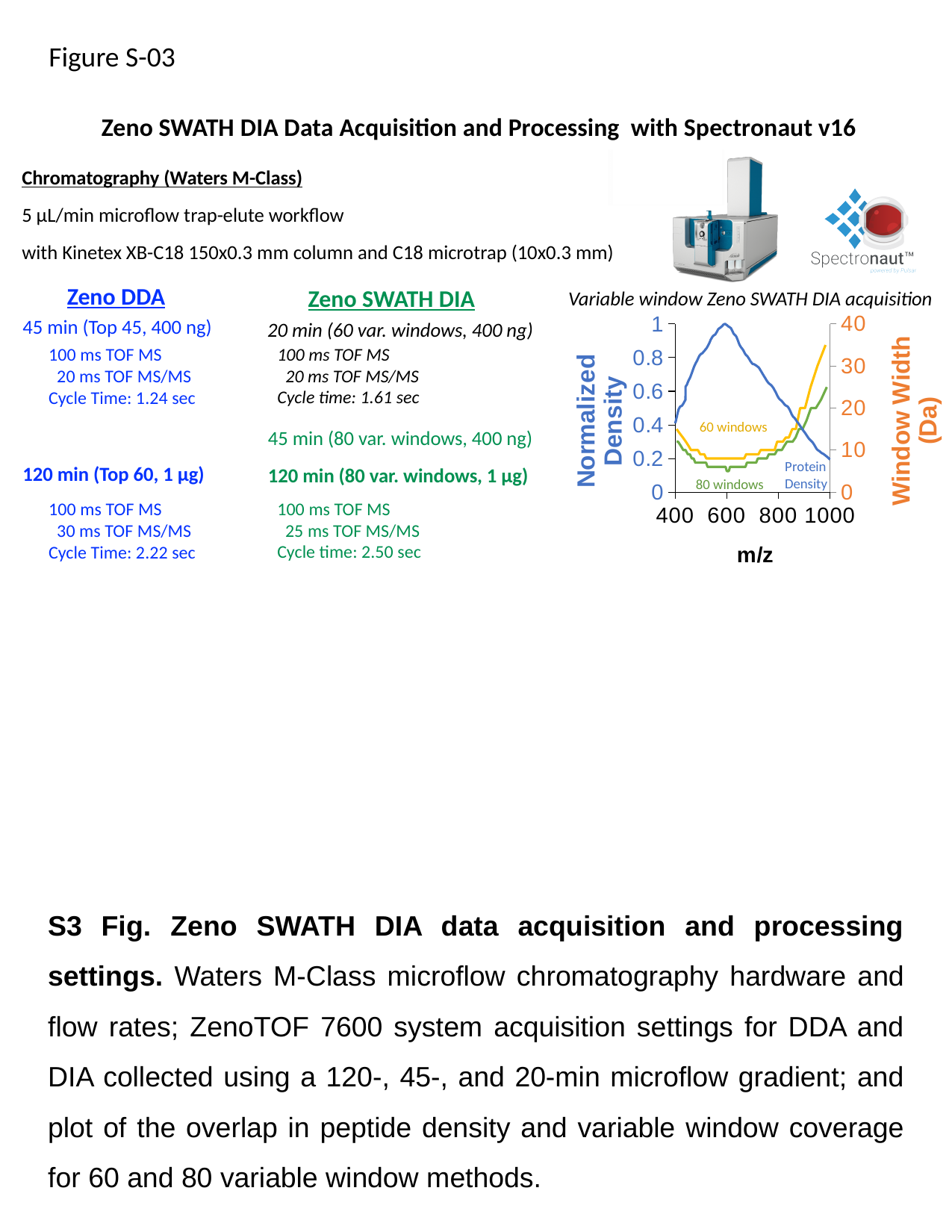

Figure S-03
#### Zeno SWATH DIA Data Acquisition and Processing with Spectronaut v16
Chromatography (Waters M-Class)
5 µL/min microflow trap-elute workflow
with Kinetex XB-C18 150x0.3 mm column and C18 microtrap (10x0.3 mm)
Zeno SWATH DIA
Zeno DDA
Variable window Zeno SWATH DIA acquisition
###### Chart
| Category | | | |
|---|---|---|---|45 min (Top 45, 400 ng)
20 min (60 var. windows, 400 ng)
100 ms TOF MS
 20 ms TOF MS/MS
Cycle time: 1.61 sec
100 ms TOF MS
 20 ms TOF MS/MS
Cycle Time: 1.24 sec
60 windows
45 min (80 var. windows, 400 ng)
Protein
Density
120 min (80 var. windows, 1 µg)
120 min (Top 60, 1 µg)
80 windows
100 ms TOF MS
 25 ms TOF MS/MS
Cycle time: 2.50 sec
100 ms TOF MS
 30 ms TOF MS/MS
Cycle Time: 2.22 sec
S3 Fig. Zeno SWATH DIA data acquisition and processing settings. Waters M-Class microflow chromatography hardware and flow rates; ZenoTOF 7600 system acquisition settings for DDA and DIA collected using a 120-, 45-, and 20-min microflow gradient; and plot of the overlap in peptide density and variable window coverage for 60 and 80 variable window methods.
