## Supplementary material for "Substantial Downregulation of Mitochondrial and Peroxisomal Proteins during Acute Kidney Injury revealed by Data-Independent Acquisition Proteomics": Suppl Figures: Figure_S-04_Zeno_Kidney_23_0224_v03.pptx

### Slide 1
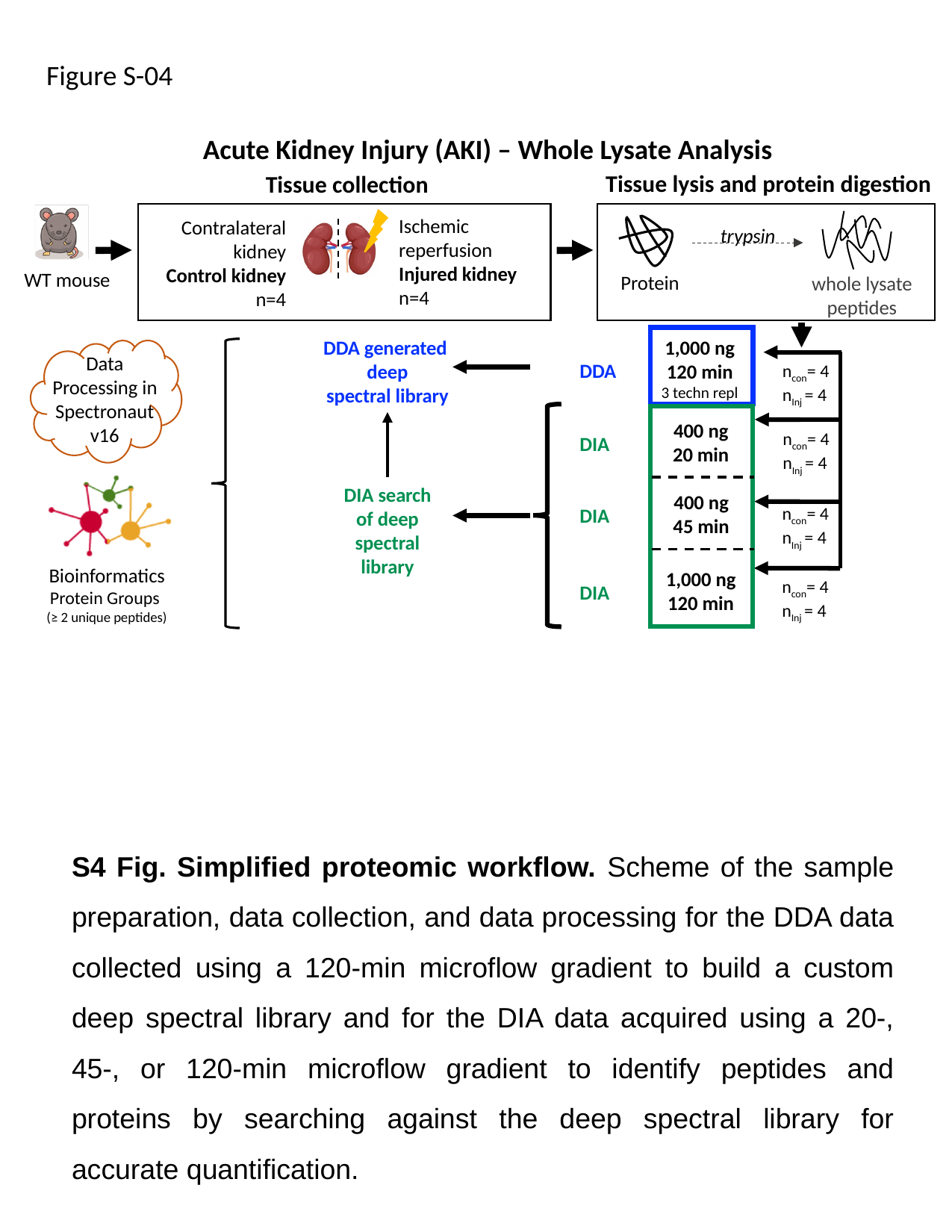

Figure S-04
Acute Kidney Injury (AKI) – Whole Lysate Analysis
Tissue lysis and protein digestion
Tissue collection
Ischemic reperfusion
Injured kidney
n=4
Contralateral kidney
Control kidney
n=4
trypsin
WT mouse
Protein
whole lysate peptides
1,000 ng
120 min
3 techn repl
DDA generated
deep
spectral library
Data Processing in Spectronaut v16
DDA
n_con_= 4
n_Inj_ = 4
400 ng
20 min
n_con_= 4
n_Inj_ = 4
DIA
DIA search
of deep
spectral library
400 ng
45 min
n_con_= 4
n_Inj_ = 4
DIA
Bioinformatics
Protein Groups
(≥ 2 unique peptides)
1,000 ng
120 min
n_con_= 4
n_Inj_ = 4
DIA
S4 Fig. Simplified proteomic workflow. Scheme of the sample preparation, data collection, and data processing for the DDA data collected using a 120-min microflow gradient to build a custom deep spectral library and for the DIA data acquired using a 20-, 45-, or 120-min microflow gradient to identify peptides and proteins by searching against the deep spectral library for accurate quantification.
