## Supplementary material for "Substantial Downregulation of Mitochondrial and Peroxisomal Proteins during Acute Kidney Injury revealed by Data-Independent Acquisition Proteomics": Suppl Figures: Figure_S-05_Zeno_Kidney_23_0224_v03.pptx

### Slide 1
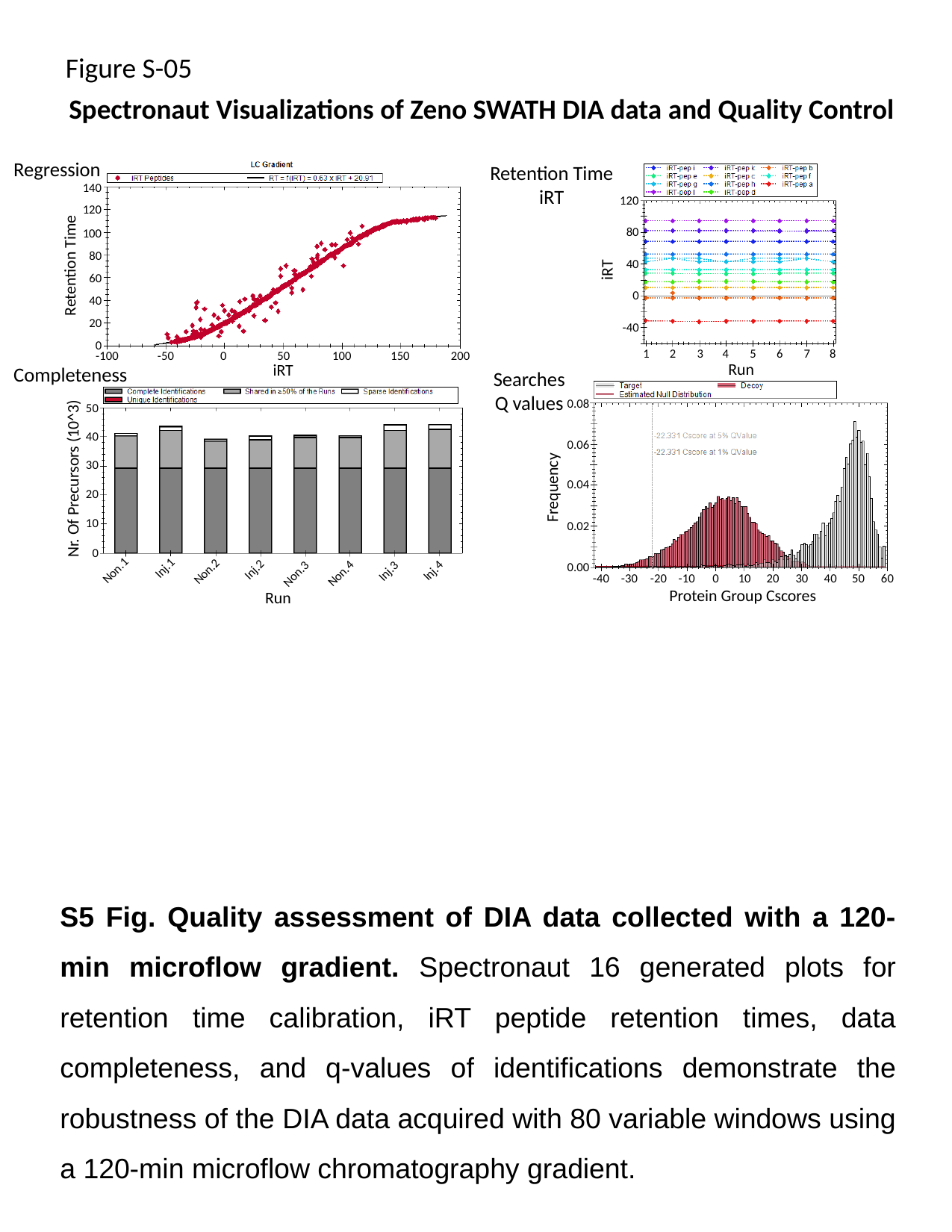

Figure S-05
Spectronaut Visualizations of Zeno SWATH DIA data and Quality Control
Regression
Retention Time
iRT
140
120
120
80
100
80
Retention Time
40
iRT
60
0
40
20
-40
0
1
2
3
4
5
6
7
8
Run
-100
-50
0
50
100
150
200
iRT
Completeness
Searches
Q values
0.08
50
40
0.06
30
Nr. Of Precursors (10^3)
0.04
Frequency
20
10
0.02
0
0.00
Non.1
Inj.1
Non.2
Inj.2
Non.4
Inj.4
Inj.3
Non.3
-40
-30
-20
-10
0
10
20
30
Protein Group Cscores
40
50
60
Run
S5 Fig. Quality assessment of DIA data collected with a 120-min microflow gradient. Spectronaut 16 generated plots for retention time calibration, iRT peptide retention times, data completeness, and q-values of identifications demonstrate the robustness of the DIA data acquired with 80 variable windows using a 120-min microflow chromatography gradient.
