## Supplementary material for "Substantial Downregulation of Mitochondrial and Peroxisomal Proteins during Acute Kidney Injury revealed by Data-Independent Acquisition Proteomics": Suppl Tables: Table_S2_Top15_Up_0224_2023_v04.docx

**Supplementary Table 23A.** Displaying the top 15 upregulated proteins, sorted by q-value, in the injured kidney.

| Proteins | Protein Description | Average Log_2_ Ratio | Q-value |
| --- | --- | --- | --- |
| Tpm1 | Tropomyosin alpha-1 chain | 1.29 | 6.15 e-12 |
| Mapre1 | Microtubule-associated protein RP/EB family member 1 | 1.07 | 7.35 e-12 |
| Ncl | Nucleolin | 1.42 | 1.57 e-10 |
| Rpl6 | 60S ribosomal protein L6 | 0.68 | 2.07 e-10 |
| Ddx3x | ATP-dependent RNA helicase DDX3X | 0.81 | 5.29 e-10 |
| Anp32a | Acidic leucine-rich nuclear phosphoprotein 32 family member A | 0.84 | 7.31 e-10 |
| Hmgb1 | High mobility group protein B1 | 0.98 | 1.02 e-09 |
| Lmna | Prelamin-A/C | 1.37 | 1.02 e-09 |
| Rplp0 | 60S acidic ribosomal protein P0 | 0.60 | 1.60 e-09 |
| Celf1 | CUG triplet repeat, RNA binding protein 1 | 1.25 | 1.77 e-09 |
| Hnrnpc | Heterogeneous nuclear ribonucleoproteins C1/C2 | 1.19 | 1.96 e-09 |
| Ppt1 | Palmitoyl-protein thioesterase 1 | 1.21 | 2.46 e-09 |
| Rpl21 | 60S ribosomal protein L21 | 0.63 | 2.49 e-09 |
| Coro1c | Coronin-1C | 1.04 | 2.49 e-09 |
| Eef1d | Elongation factor 1-delta | 0.67 | 2.65 e-09 |

**Supplementary Table 2B.** Displaying the top 15 up-regulated proteins in injured kidney sorted by log_2_(FC) from the 120 min microflow gradient DIA data.

| Proteins | Protein Description | | Average Log_2_ Ratio | Q-value |
| --- | --- | --- | --- | --- |
| Havcr1 | | Hepatitis A virus cellular receptor 1 homolog | 7.13 | 1.42 e-06 |
| Vtn | | Vitronectin | 5.27 | 5.30 e-04 |
| Mup3 | | MCG15829 | 5.14 | 3.89 e-06 |
| Serpina3n | | Serine (Or cysteine) peptidase inhibitor | 5.05 | 4.04 e-04 |
| C1qb | | Complement C1q subcomponent subunit B | 4.85 | 1.73 e-03 |
| Lcn2 | | Neutrophil gelatinase-associated lipocalin | 4.76 | 3.14 e-06 |
| Apon | | Apolipoprotein N | 4.53 | 2.23 e-03 |
| Mup17 | | Major urinary protein 17 | 4.35 | 5.04 e-06 |
| C1qc | | Complement C1q subcomponent subunit C | 4.33 | 1.70 e-03 |
| Mup14 | | Protein Mup14 (Fragment) | 4.26 | 2.78 e-06 |
| Thy1 | | Thy-1 membrane glycoprotein | 4.21 | 2.47 e-05 |
| Hspb1 | | Heat shock protein beta-1 | 4.19 | 2.04 e-06 |
| Mt2 | | Metallothionein-2 | 4.18 | 6.94 e-06 |
| Ngp | | Neutrophilic granule protein | 4.11 | 1.70 e-03 |
| Mt1 | | Metallothionein-1 | 3.97 | 3.63 e-06 |
