## Supplementary material for "Substantial Downregulation of Mitochondrial and Peroxisomal Proteins during Acute Kidney Injury revealed by Data-Independent Acquisition Proteomics": Suppl Tables: Table_S3_Top15_Down_0224_2023_v04.docx

**Supplementary Table 3A.** Displaying the top 15 downregulated proteins, sorted by q-value, in the injured kidney.

| Proteins | Protein Descriptions | Subcellular Location | Average Log_2_ Ratio | Q-value |
| --- | --- | --- | --- | --- |
| Acox2 | Peroxisomal acyl-coenzyme A oxidase 2 | Peroxisome | -2.46 | 7.93 e-12 |
| Amacr | Alpha-methylacyl-CoA racemase | Mitochondria, Peroxisome | -2.21 | 2.84 e-11 |
| Idh2 | Isocitrate dehydrogenase [NADP], mitochondrial | Mitochondria | -1.55 | 2.84 e-11 |
| Slc34a1 | Sodium-dependent phosphate transport protein 2A | Cell Membrane | -2.83 | 4.41 e-11 |
| Hsd17b8 | Estradiol 17-beta-dehydrogenase 8 | Mitochondria | -1.58 | 8.03 e-11 |
| Dmgdh | Dimethylglycine dehydrogenase, mitochondrial | Mitochondria | -1.74 | 9.46 e-11 |
| Pah | Phenylalanine-4-hydroxylase | Cytosol | -2.38 | 1.47 e-10 |
| Mccc2 | Methylcrotonoyl-CoA carboxylase beta chain, mitoch. | Mitochondria | -1.61 | 1.57 e-10 |
| Ndufs6 | NADH dehydrogenase iron-sulfur protein 6, mitoch. | Mitochondria | -1.51 | 1.57 e-10 |
| Opa1 | Isoform 2 of Dynamin-like 120 kDa protein, mitochondrial | Mitochondria | -1.24 | 1.57 e-10 |
| Kyat3 | Kynurenine--oxoglutarate transaminase 3 | Cytosol, Mitochondria | -2.45 | 1.91 e-10 |
| Akr1c14 | 3-alpha-hydroxysteroid dehydrogenase type 1 | Cytosol | -2.94 | 2.17 e-10 |
| Hint2 | Histidine triad nucleotide-binding protein 2, mitochondrial | Mitochondria | -1.54 | 2.17 e-10 |
| Idh3a | Isocitrate dehydrogenase [NAD] subunit alpha, mitochondrial | Mitochondria | -1.41 | 2.17 e-10 |
| Hadha | Trifunctional enzyme subunit alpha, mitochondrial | Mitochondria | -1.14 | 2.17 e-10 |

**Supplementary Table 3B.** Displaying the top 15 down-regulated proteins in injured kidney sorted by log_2_(FC) from the 120 min microflow gradient DIA data.

| Proteins | Protein Descriptions | Average Log_2_ Ratio | Q-value |
| --- | --- | --- | --- |
| Uroc1 | Urocanate hydratase | -3.62 | 6.09 e-09 |
| Ppp1r1a | Protein phosphatase 1 regulatory subunit 1A | -3.33 | 2.71 e-08 |
| Bsnd | Barttin | -3.20 | 2.67 e-07 |
| Bcat1 | Branched-chain-amino-acid aminotransferase, cytosolic | -3.19 | 1.45 e-06 |
| Mpv17l | Mpv17-like protein | -3.16 | 4.18 e-04 |
| Akr1c14 | 3-alpha-hydroxysteroid dehydrogenase type 1 | -2.94 | 2.17 e-10 |
| Nccrp1 | F-box only protein 50 | -2.86 | 4.90 e-07 |
| Slc34a1 | Sodium-dependent phosphate transport protein 2A | -2.83 | 4.41 e-11 |
| Prkaa2 | 5'-AMP-activated protein kinase catalytic subunit alpha-2 | -2.83 | 2.34 e -04 |
| Dnajc12 | DnaJ homolog subfamily C member 12 | -2.82 | 8.98 e-09 |
| Msrb2 | Methionine-R-sulfoxide reductase B2, mitochondrial | -2.69 | 5.10 e-09 |
| Slc12a1 | Solute carrier family 12 member 1 | -2.65 | 1.79 e-08 |
| Ca14 | Carbonic anhydrase 14 | -2.62 | 1.68 e-07 |
| Slc22a22 | BC026439 protein | -2.60 | 5.23 e-09 |
| Cyp2d9 | Cytochrome P450 2D9 | -2.60 | 4.08 e-05 |
